# Abelson kinase’s intrinsically disordered linker plays important roles in protein function and protein stability

**DOI:** 10.1101/2020.05.20.106708

**Authors:** Edward M. Rogers, S. Colby Allred, Mark Peifer

**Affiliations:** Department of Biology, University of North Carolina at Chapel Hill, Chapel Hill, NC 27599, USA; Lineberger Comprehensive Cancer Center, University of North Carolina at Chapel Hill, Chapel Hill, NC 27599, USA

## Abstract

The non-receptor tyrosine kinase Abelson (Abl) is a key player in oncogenesis, with kinase inhibitors serving as paradigms of targeted therapy. Abl also is a critical regulator of normal development, playing conserved roles in regulating cell behavior, brain development and morphogenesis. Drosophila offers a superb model for studying Abl’s normal function, because, unlike mammals, there is only a single fly Abl family member. Abl has multiple roles in embryonic morphogenesis, and we and others have begun to take Abl apart as a machine. This revealed many surprises. For instance, kinase activity, while important, is not crucial for all Abl activities, and the C-terminal F-actin binding domain plays a very modest role. This turned our attention to less well-known feature—the long intrinsically-disordered region (IDR) linking Abl’s kinase and F-actin binding domains. The past decade revealed unexpected, important roles for IDRs in diverse cell functions, by mediating multivalent interactions, enabling assembly of biomolecular condensates via phase separation. Previous work deleting conserved regions revealed an important role for a PXXP motif in the IDR, but did not identify any other essential regions. Here we extend this, deleting the entire IDR. This revealed essential roles for the IDR in embryonic and adult viability, and in cell shape changes and cytoskeletal regulation during embryonic morphogenesis. Strikingly, AblΔIDR acts as dominant negative, worsening the phenotype of the null mutant. AblΔIDR accumulates at >10-fold higher levels than wildtype Abl in both Drosophila embryos and cultured cells, suggesting important roles for the IDR in modulating protein stability.

## Introduction

Biomedical research has dual goals: to uncover mechanisms underlying normal cellular function and to apply this understanding to develop better treatments in human disease. Perhaps no story better illustrates this than the discovery more than 60 years ago of the “Philadelphia chromosome”, a translocation between chromosomes 9 and 22 present only in leukocytes from patients with chronic myelogenous leukemia. It provided the first molecular link between genetics and cancer, and ultimately led to the realization that Abelson kinase (Abl) is the initiating oncogene in many cases of chronic myelogenous and acute lymphoblastic leukemia (Ren 2005). These translocations fuse the *Bcr* and *Abl* genes, removing a myristoylation sequence at Abl’s N-terminus that inhibits kinase activation, rendering the kinase constitutively active. Drugs targeting Abl kinase activity like Gleevec (Imatinib) have emerged as paradigms of targeted therapy (Sawyers *et al*. 2002; Talpaz *et al*. 2002), and spurred development of similar inhibitors of other oncogenic kinases.

Of course, non-receptor tyrosine kinases like Src and Abl kinases did not evolve to cause cancer. Both play key roles in signal transduction, regulating embryonic development and tissue homeostasis. Abl family members regulate morphogenetic movements during embryogenesis in both mammals and Drosophila, and also play key roles in neural development, axon outgrowth, and synaptogenesis (reviewed in (Moresco and Koleske 2003; Bradley and Koleske 2009; Khatri *et al*. 2016; Kannan and Giniger 2017). They act downstream of diverse receptors, including receptor tyrosine kinases or receptor serine/threonine kinases, as well as the cell-matrix and cell-cell adhesion receptors, integrins and cadherins. Downstream, Abl family members activate cytoskeletal effectors to directly regulate cell behavior, though transcriptional effectors are also important.

Abl’s structure facilitates the link between cell signaling and cytoskeletal regulation. All Abl family members share a highly conserved set of N-terminal domains with Src (Fig. 1A). These include a Src homology 2 (SH2) domain that binds specific peptides carrying a phosphorylated tyrosine and an SH3 domain that binds specific proline-rich peptides, both allowing interactions with upstream receptors and downstream effectors. These are immediately followed by the conserved tyrosine kinase domain (Colicelli 2010). However, unlike Src, Abl family members have long C-terminal extensions, with a C-terminal F-actin-binding domain (FABD) separated from the N-terminal module by a long, poorly conserved linker that is predicted to be intrinsically disordered (the intrinsically disordered region or IDR). Different family members share peptide motifs within the IDR that bind or are predicted to bind actin, microtubules, Ena/VASP family members, and SH3 domain containing proteins. The only peptide motif in the IDR clearly conserved between mammals and *Drosophila* is a PXXP motif that in mammals binds both SH2/SH3 adapters like Crk and NCK and the actin regulator Abl interacting protein (Abi) (Hossain *et al*. 2012; Gregor *et al*. 2019).

**Figure 1.**
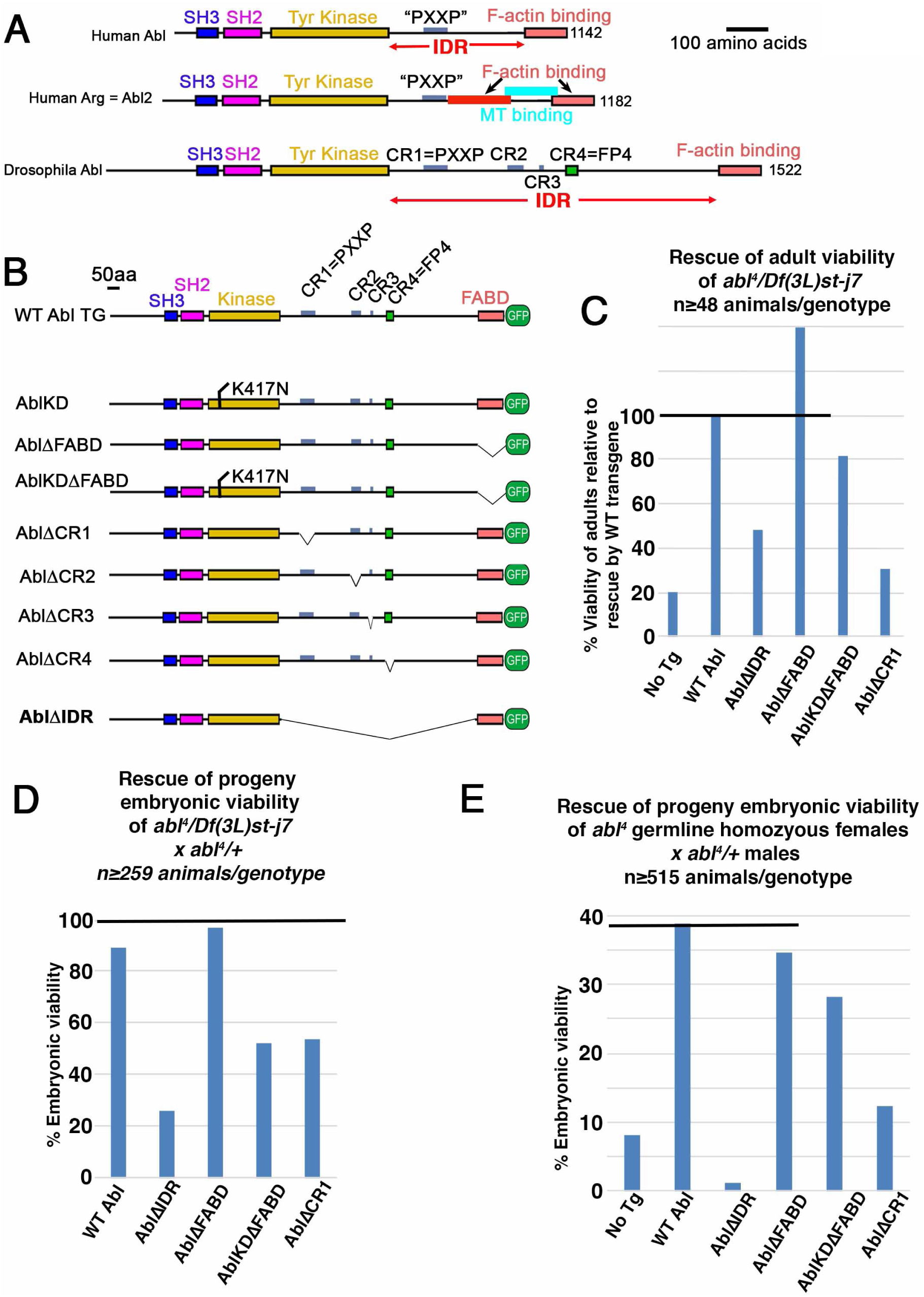
Generating AblΔIDR and testing its ability to rescue adult and embryonic viability. A. Diagram of human Abl and Arg and Drosophila Abl, showing conserved domains/motifs as well as motifs in the IDR that vary between family members. B. Illustration of the mutant Abl proteins we previously tested and our new AblΔIDR mutant. It was designed to remove essentially the entire IDR, leaving only a few amino acids at each end to ensure we did not disrupt folding of the kinase domain or FABD. A 15 aa flexible linker was added in its place. C. Assessment of the ability of AblΔIDR to rescue the viability of *abl*^*4*^*/DfAbl* adults, normalized to rescue by our wildtype Abl transgene, and compared to rescue by some of our previously tested mutants. Full data sets with statistical significance for C-E are in Table 1. D. Assessment of the ability of AblΔIDR to rescue embryonic viability of the progeny of *abl*^*4*^*/DfAbl* females mated to *abl*^*4*^*/+* males, compared to rescue by some of our previously tested mutants. Line indicates 100% embryonic viability. E. Assessment of the ability of AblΔIDR to rescue embryonic viability of the progeny females with germlines homozygous for of *abl*^*4*^ mated to *abl*^*4*^*/+* males, compared to rescue by some of our previously tested mutants. Line indicates degree of rescue by our wildtype Abl transgene.

Mammals have two Abl family members, Abl and Abl-related gene (Arg), with partially redundant functions in development and tissue homeostasis. *Abl* single mutant mice die neonatally with thymic and splenic atrophy, T and B cell lymphopenia, osteoporosis, and cardiac defects (Schwartzberg *et al*. 1991; Tybulewicz *et al*. 1991; Hardin *et al*. 1995; Hardin *et al*. 1996; Li *et al*. 2000; Qiu *et al*. 2010b). Conditional knockout confirmed roles in T cells (Zipfel *et al*. 2004; Gu *et al*. 2007; Huang *et al*. 2008). *Arg* single mutant mice are viable with grossly normal brains, but exhibit multiple behavioral defects (Koleske *et al*. 1998), likely linked to a reduced ability to maintain dendrites (Lin *et al*. 2013). Arg mutants also exhibit subtle defects in muscle development (Lee *et al*. 2017). In contrast, loss of both Abl and Arg leads to embryonic lethality at day 11, with a failure to complete neural tube closure. Conditional double knockout has revealed additional important redundant roles in cerebellar (Qiu *et al*. 2010a), cerebral (Moresco *et al*. 2005), and endothelial development and barrier function (Chislock and Pendergast 2013; Chislock *et al*. 2013).

**Table 1.**

| Construct | Residues changed or deleted | Percent rescue of adult viability in *abl*<sup>4</sup> / *Df(3L)st-j7* (normalized to WT) | Probabability that viability is **higher** than than adults with no transgene | Probabability that rescue is **better** or **worse** than that by Abl WT | Percent rescue of progeny embryonic viability of *abl*<sup>4</sup> / *Df(3L)st-j7* X *abl*<sup>4</sup> / + | Probabability that rescue is worse than that by Abl WT | Percent rescue of progeny embryonic viability of *abl*<sup>4</sup> maternal germ line clone X *abl4* / + | Probabability that lethality is **worse** than that of Abl WT | Probabability that lethality is **better** or **worse** than that of *abl* / *MZ* |
| --- | --- | --- | --- | --- | --- | --- | --- | --- | --- |
| No Transgene | N/A | 20.4 (n=145) | N/A | **p<0.0001** | N/A | N/A | 8.2% (n=527) # | **p<0.0001** | n/a |
| Abl WT | N/A | 100% (n=386) | **p<0.0001** | N/A | 89.4 (n=901) # | N/A | 38.9% (n=540) # | n/a | **p<0.0001** |
| AblΔIDR | 679-1398 | 48.1% (n=223) | **p=0.0075** | **p<0.0001** | 26.2% (n=378) | p<0.0001 | 1.1% (n=569) | **p<0.0001** | **p<0.0001** |
| AblΔFABD | 1418-1520 | 140% (n=173) | **p<0.0001** | **p<0.0001** | 97.1 (n=362) # | N.S. | 34.6% (n=515) # | p=0.16 | **p<0.0001** |
| AblKDΔFABD | K417N,1418-1520 | 81.5% (n=585) | **p<0.0001** | **p=0.0108** | 52.1 (n=259) # | p<0.0001 | 28.2 (n=560) # | **p=0.0002** | **p<0.0001** |
| AblΔCR1 | 734-789 | 30.7% (n=48) | p=0.4474 | **p<0.0001** | 53.9 (n=267) # | p<0.0001 | 12.4% (n=606) # | **p<0.0001** | **p=0.02** |

n=number of adults/embryos scored; N/A=not applicable; NS=not significant Green text=better than no transgene or AblWT Red text =worse than no transgene or AblWT

*Drosophila* has a single Abl family member, simplifying analysis of its roles in development. In the 1980s Michael Hoffmann and his lab identified the first mutations in fly Abl, as part of a pioneering effort to define the normal roles of human oncogenes (Henkemeyer *et al*. 1987). Others built on these early efforts. Like its mammalian homologs, Abl plays important roles in embryonic and postembryonic neural development, with Abl acting downstream of diverse axon guidance receptors, including DCC/Frazzled, Robo, Plexin, Eph, and Notch (reviewed in (Kannan and Giniger 2017). Genetic and cell biological analyses also revealed Abl’s downstream effectors, the most prominent of which is Enabled (Ena), which binds the growing end of actin filaments and promotes their elongation. Abl negatively regulates Ena (Gertler *et al*. 1990), through a mechanism that remains unclear. Trio, a GTP exchange factor (GEF) for the small GTPase Rac, is also an Abl effector.

Subsequent analysis of embryos lacking both maternal and zygotic Abl revealed additional roles outside of the nervous system. Abl regulates diverse events ranging from the actin-dependent cellularization process, to apical constriction of mesoderm precursors, cell intercalation during germband elongation, and collective cell migration during germband retraction, dorsal closure, and head involution (Grevengoed *et al*. 2001; Grevengoed *et al*. 2003; Fox and Peifer 2007; Tamada *et al*. 2012). In these events, regulation by and of the cadherin-based cell adhesion machinery plays a role, while Ena remains a critical downstream target (Grevengoed *et al*. 2001; Grevengoed *et al*. 2003; Fox and Peifer 2007).

Abl kinase activity has been a focus of much attention, particularly after the success of Abl kinase inhibitors in the treatment of leukemia. However, the simplistic picture of Abl as a kinase acting solely by phosphorylating downstream proteins rapidly proved inaccurate. Kinase-dead Abl rescues defects in both adult viability and retinal development (Henkemeyer *et al*. 1990). Analysis of its role in embryonic development suggests kinase activity is important for roles in both axon patterning and in morphogenesis, but a kinase-dead mutant retains significant residual function (O’donnell and Bashaw 2013; Rogers *et al*. 2016). More limited analysis implicated the SH2 domain in axon guidance (O’donnell and Bashaw 2013). The extended C-terminal region of Abl, including both the IDR and the C-terminal FABD, is essential for function, as *abl*^*1*^, which encodes a stable protein truncated soon after the kinase domain, behaves genetically as a null allele (Henkemeyer *et al*. 1987). Similar results were seen in mice where a truncated protein resembled the null (Schwartzberg *et al*. 1991; Tybulewicz *et al*. 1991). The simplest explanation would be that this reflected an essential function of the FABD. However, surprisingly Abl lacking the FABD fully rescues viability and fertility, though detailed analysis of axon patterning and synergistic effects with loss of kinase activity suggest the FABD does play a supporting role (O’donnell and Bashaw 2013; Rogers *et al*. 2016; Cheong and Vanberkum 2017).

These data opened up potential roles for the IDR. The past decade revealed unexpected and important roles for IDRs in diverse cell functions (Oldfield and Dunker 2014), through their ability to mediate multivalent interactions, including those enabling assembly of “biomolecular condensates”. These condensates organize proteins and RNAs into non-membrane bound cellular compartments that perform diverse functions, ranging from regulating transcription to RNA processing/RNP assembly to the DNA damage response to cellular signaling (Banani *et al*. 2017; Holehouse and Pappu 2018). Biomolecular condensates assemble by multivalent interactions among their protein and RNA components, leading to “phase separation”. While not all proteins containing IDRs have been shown to form biomolecular condensates, intriguingly proteins containing SH2 and SH3 domains were among the first proteins shown to assemble by this mechanism (Li *et al*. 2012), and Abl clearly can assemble into a large macromolecular complex (e.g. (Gregor *et al*. 2019).

Abl’s IDR is not highly conserved in primary sequence, even between the two mammalian paralogs (Fig. 1A). Only a single peptide is conserved among fly and mammalian family members—the PXXP SH3-domain binding motif in the N-terminal quarter of the IDR. Other peptide motifs, including a predicted Ena binding site, are conserved over shorter phylogenetic distances; e.g., among insect Abl proteins. There are also functional motifs present only in single family members, including the microtubule and second actin interaction sites in mammalian Arg (Miller *et al*. 2004; Courtemanche *et al*. 2015). Two groups assessed the function of the *Drosophila* Abl IDR, taking different approaches. Our lab individually deleted short conserved regions of 12–56 amino acids (conserved regions 1 to 4 (CR1-CR4)), in the context of a GFP-tagged full length Abl protein driven by its endogenous promotor (Fig. 1B). We measured rescue of embryonic and adult viability, morphogenetic movements in the embryo, and axon outgrowth in the embryonic central nervous system (Rogers *et al*. 2016). Cheong and VanBerkum took a more comprehensive approach, deleting successively smaller fractions of the C-terminal region, including the FABD. They began by dividing it in half, then in quarters and then focused in on two smaller regions, with smaller deletions and point mutations. They expressed their mutant proteins in the background of zygotic *abl* mutants using the GAL4-UAS system and assessed rescue of axon pathfinding (Cheong and Vanberkum 2017). Both approaches led to similar surprising conclusions. Only a single region of the IDR plays a major role in Abl function—the region containing the conserved PXXP motif. Surprisingly, however, this motif was extremely important, as its deletion caused as reduced Abl function more than loss of kinase activity or even loss of both kinase activity and the FABD. Subsequent analyses support the idea that this motif acts by interactions with the adapter protein Crk (Spracklen *et al*. 2019) and with the actin-regulatory WAVE-regulatory complex (Cheong *et al*. 2020). The fine-grained dissections of the IDR by Cheong and VanBerkum suggest other regions of the IDR may have more subtle roles in axon guidance.

Our initial analyses did not fully probe the function of the IDR, as perhaps the simplest test of its function—completely deleting the IDR while leaving the FABD intact—was missed. We thus generated a mutation cleanly removing the IDR in the context of a GFP-tagged Abl construct driven by its endogenous promotor. This revealed surprising dominant-negative activity of AblΔIDR and a potential role for the IDR in regulation of Abl protein stability. These observations provide important insights into Abl function.

## Results

### Creating a mutant to test the role of the IDR in Abl function

Abl is a multidomain protein which uses both its kinase activity and its protein interaction domains to create a signaling hub, integrating upstream signals and activating downstream effectors. Our lab previously created a series of *abl* mutants to assess the role of kinase activity and other domains and motifs in *Drosophila* (Fig. 1B; (Rogers *et al*. 2016). The base construct was a P-element transgene containing a wild type *abl* gene driven by a 2kb fragment of the 5’ upstream endogenous *abl* promoter (Tn Abl WT:GFP), which can fully rescue *ablMZ* mutant embryos (Fox and Peifer 2007). This construct has a C-terminal GFP tag that does not impair its rescuing ability, and which allows direct visualization of Abl localization in live embryos. These included mutants deleting short conserved regions in the IDR (AblΔCR1-ΔCR4,Fig 1B), but we did not fully remove the IDR to assess its full set of roles. To do so, here we created a similar transgene that essentially deletes the entire IDR—below we refer to this as AblΔIDR (Fig. 1B; amino acids 679-1398 are deleted; see Methods for details). We added a 15 amino acid flexible linker (GGS)_5_) in place of the IDR to reduce the likelihood of disrupting folding of the adjacent kinase domain and FABD (Van Rosmalen *et al*. 2017). We introduced this transgene into the *Drosophila* genome in two ways—by P element-based transformation (selecting an insertion on then second chromosome), and by site-specific integration on the left arm of the 2nd chromosome (at 22A3). We then used these transgenes to assess the roles of the IDR in Abl function.

### AblΔIDR does not rescue adult viability and has apparent dominant negative effects on embryonic morphogenesis

Because of its critical roles in embryonic morphogenesis and neuronal development, Abl is essential for both embryonic and adult viability (Henkemeyer *et al*. 1987; Grevengoed *et al*. 2001). The ability to rescue viability thus offered an initial test for our AblΔIDR mutant protein. Abl is maternally contributed and this maternal contribution is sufficient for embryonic development (Grevengoed *et al*. 2001). However, most *abl* null mutants die as pupae—the few that escape are functionally sterile and die soon after eclosing (Henkemeyer *et al*. 1987). We thus tested the ability of AblΔIDR to rescue adults that were heterozygous for the putative null allele *abl*^*4*^ (Fox and Peifer 2007) and a Deficiency, *Df(3L)st-7 Ki*. that removes the *abl* gene (*abl*^*4*^*/Df*), as we had for our earlier mutants (Rogers *et al*. 2016). A targeted transgene encoding AblΔIDR provided partial but incomplete rescue of adult viability. While unrescued *abl*^*4*^*/Df* adults had only 20% the viability of those rescued by our wildtype *abl* transgene, AblΔIDR; *abl*^*4*^*/Df* adults eclosed at 48% the rate of those rescued by the wildtype transgene (Fig.1C;Table 1). In contrast, AblΔFABD fully rescued adult viability, and even a mutant lacking both kinase activity and the FABD provided substantial rescue (Rogers *et al*. 2016); 82% of the wildtype transgene). However, AblΔCR1, lacking the PXXP motif in the IDR, did not provide substantial rescue (31% viability relative to the wildtype transgene; Fig. 1C; Table 1; (Rogers *et al*. 2016). These data suggest that the IDR is important for Abl function.

Unlike the *abl*^*4*^*/Df* escapers, AblΔIDR; *abl*^*4*^*/Df* females lived long enough to mate and produce fertilized eggs. We thus asked whether AblΔIDR rescued the lethality of embryos lacking both maternal and zygotic Abl, by crossing these females to males who were heterozygous for *abl*^*4*^*/+* and carried the transgene. AblΔIDR did not rescue the viability of maternal/zygotic mutants (50% of the progeny), and, surprisingly, even 30% of embryos that inherited a paternal zygotic wildtype *abl* gene died before hatching and the rest (20%) died as first instar larvae (Fig. 1D, Table 1). In contrast, AblΔFABD provided full rescue of embryonic viability (Fig. 1D; Table 1; (Rogers *et al*. 2016). To roughly assess the rescue of embryonic morphogenesis by AblΔIDR, we examined the cuticles of the dead embryos. To our surprise, the cuticle phenotype was <u>extremely</u> severe, with all embryos exhibiting strong disruption of epidermal integrity, including those in which only fragments of cuticle were secreted (Fig. 2A vs B-D). These morphogenetic phenotypes are more severe than those characteristically seen in *abl* maternal/zygotic mutant embryos (Grevengoed *et al*. 2001; Rogers *et al*. 2016). However, the limitations of this approach are that since unrescued *abl*^*4*^*/Df* mutant females are sterile, we could not compare embryonic morphogenesis of their progeny to those rescued by AblΔIDR.

**Figure 2.**
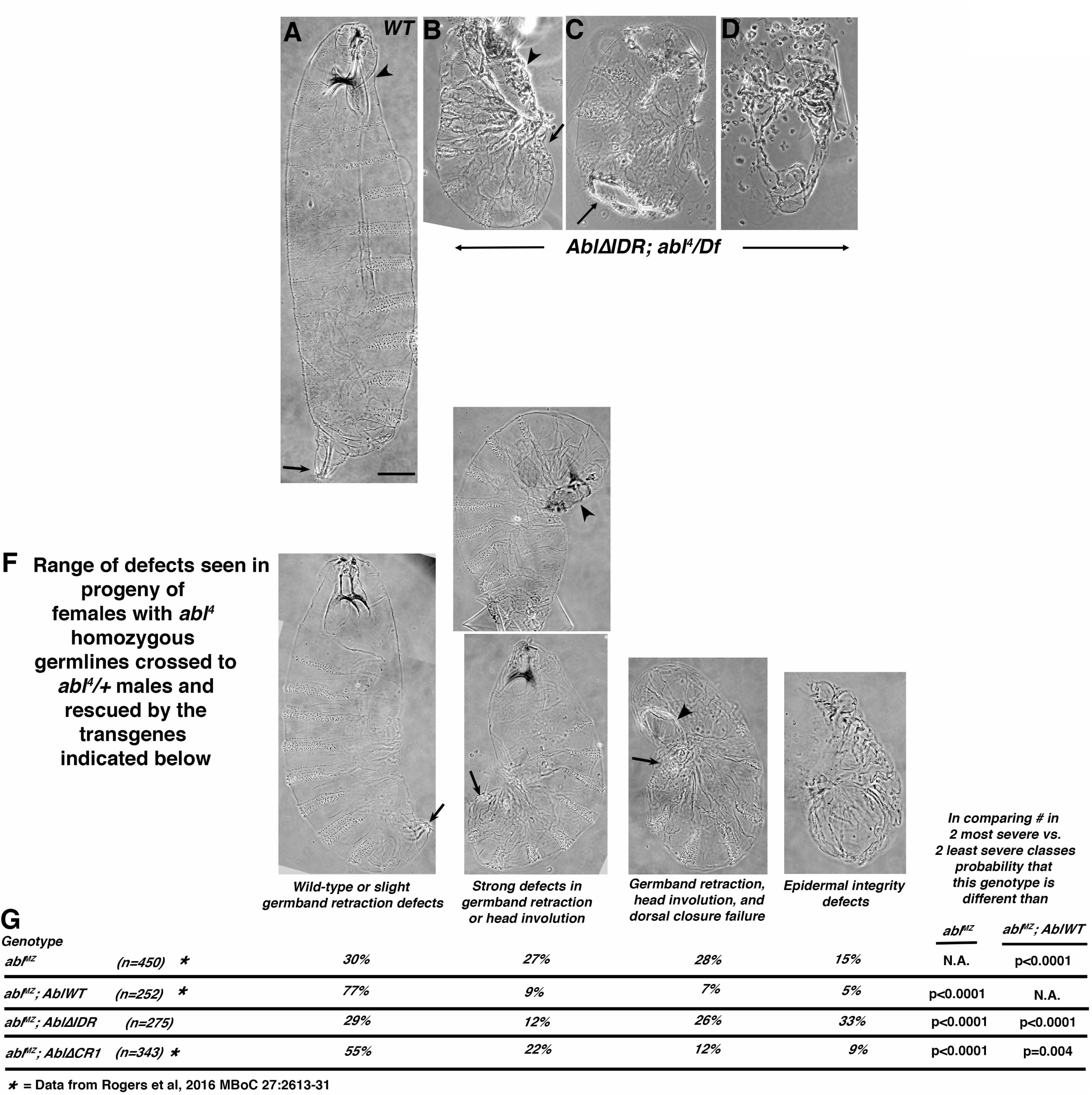
AblΔIDR does not rescue embryonic morphogenesis. A-F. Cuticle preparations. Anterior up. A. Wildtype, ventral side right, revealing the segmental array of denticle belts and naked cuticle. Arrowhead: head involution was completed and there is a well-formed head skeleton. Arrow. Germband retraction was completed, positioning the spiracles at the posterior end. Scale bar=50 µm. B-D. Examples of cuticles from progeny of *AblΔIDR; abl*^*4*^*/Df* mothers crossed to *abl*^*4*^*/+* fathers. B. Least severe phenotype. Head involution, dorsal closure (arrowhead) and germband retraction (arrow) failed. C. Intermediate phenotype, with large hole in the ventral cuticle. D. Severe phenotype. Only fragments of cuticle remain. F. Range of cuticle defects seen in the progeny of females whose germlines are homozygous for *abl*^*4*^ crossed to *abl*^*4*^*/+* fathers, carrying the transgenes indicated in G maternally and zygotically. Arrows and arrowheads as in A-D. Images in A and F are from Rogers et al., 2016, where we developed this cuticle scoring scheme. G. Frequencies of each phenotype in the indicated genotypes. Statistical test used was Fisher’s Exact Test.

To circumvent this, we used the FLP/FRT/DFS approach (Chou *et al*. 1993) to generate females whose germlines are homozygous for *abl*^*4*^, either in the presence of one of our transgenes or in the absence of any transgene as a control. This approach allowed us to compare maternal/zygotic *abl* mutants (*ablMZ)*, who are homozygous for the null allele, with similar mutants that have one of our transgenes contributed both maternally and zygotically. We used *abl* transgenes inserted at the same chromosomal location via phiC integrase. *ablMZ* mutants generated by the FLP/FRT/DFS approach are embryonic lethal (Grevengoed *et al*. 2001), and there is only partial rescue of viability in the 50% of embryos that receive a wild-type *abl* gene paternally (9% overall embryonic viability (Fig. 1E; Table 1). Strikingly, AblΔIDR; *ablMZ* mutants had an <u>even higher</u> embryonic lethality (1% overall embryonic viability; probability that viability is lower than *ablMZ* p<0.0001; by Fisher’s Exact test; Table 1). In contrast, our GFP-tagged wildtype transgene, provided strong rescue (39% viability (Rogers *et al*. 2016); we attribute the lack of full rescue to other mutations that have accumulated on the *abl*^*4*^ chromosome), as did the AblΔFABD transgene (35% viability; (Rogers *et al*. 2016).

We next examined embryonic cuticles, as they allow us to assess cell fate choice, major morphogenetic movements like germband retraction, head involution, and dorsal closure, along with epidermal integrity. *ablMZ* mutants have multiple defects in these processes (Grevengoed *et al*. 2001; Rogers *et al*. 2016); Fig 2F,G), with smost exhibiting strong defects in head involution and failure of full germband retraction. Many also fail in dorsal closure, and a small fraction (15%) have defects in epidermal integrity. Our transgene encoding wildtype Abl largely rescued these defects (Fig. 2F,G; (Rogers *et al*. 2016). Our previous analysis revealed that neither kinase activity nor the FABD is essential for rescuing these cuticle defects, while AblΔCR1, lacking the conserved PXXP motif in the IDR, largely rescued epidermal integrity but only partially rescued germband retraction and dorsal closure (Fig. 2F,G; (Rogers *et al*. 2016). In contrast, however, AblΔIDR; *ablMZ* mutants had even <u>more severe</u> cuticle defects than unrescued *ablMZ* mutants. For example, the fraction of embryos with the more severe epidermal defects more than doubled, from 15% to 33% (Fig. 2F,G; probability that the cuticle defects are worse than *ablMZ* p<0.0001; by Fisher’s Exact test). This epidermal disruption phenotype was similar to that we observed in the progeny of AblΔIDR; *abl*^*4*^*/Df* females (Fig. 2B-D). Taken together, the increased embryonic lethality and higher proportion of severe cuticle defects in these experiments and the unexpectedly severe cuticle phenotype seen in our initial *abl*^*4*^*/Df* experiments, suggested to our surprise that expressing AblΔIDR not only fails to rescue loss of Abl, but actually worsens some aspects of the *abl* null mutant phenotype in embryonic development.

### AblΔIDR does not effectively rescue defects in cellularization or mesoderm invagination

Abl has diverse roles in embryonic development, ranging from regulating actin dynamics during syncytial development and cellularization to regulating apical constriction of mesodermal cells to regulating cell shape change and collective cell migration during germband retraction and dorsal closure. Our cuticle data suggested that AblΔIDR was substantially impaired in morphogenesis. To examine this more closely, we used immunofluorescence and confocal microscopy to examine cell shape changes and cytoskeletal regulation during embryonic development, as we had done to assess the roles of kinase activity, the FABD, and the conserved motifs in the IDR (Rogers *et al*. 2016).

The first events of embryogenesis requiring Abl function are the characteristic dynamics of the actin cytoskeleton during the syncytial stages and cellularization. Maternal/zygotic *abl* mutants (*ablMZ)* have defects in both processes, and thus accumulate multinucleate cells at the end of cellularization (Grevengoed *et al*. 2003); Fig. 3A vs B, red arrows). AblΔIDR did not rescue these defects, and thus AblΔIDR; *ablMZ* mutants accumulated multinucleate cells (Fig. 3A vs C, red arrows). Abl is also required for the first event of gastrulation, in which cells along the ventral midline apically constrict in a coordinated way and invaginate as a tube (Fox and Peifer 2007). The invaginating cells then go on to become mesoderm, while the ectodermal cells close the gap and form a straight midline. In *ablMZ* mutants, apical constriction is poorly coordinated, leaving some mesodermal cells on the surface. Ectodermal cells eventually close the gap, but the resulting midline is not straight (Fig.3D vs. E, blue arrows). Once again, AblΔIDR did not fully rescue these defects (Fig. 3F). This latter phenotype is particularly interesting as AblΔCR1 did rescue mesoderm invagination (Rogers *et al*. 2016).

**Figure 3.**
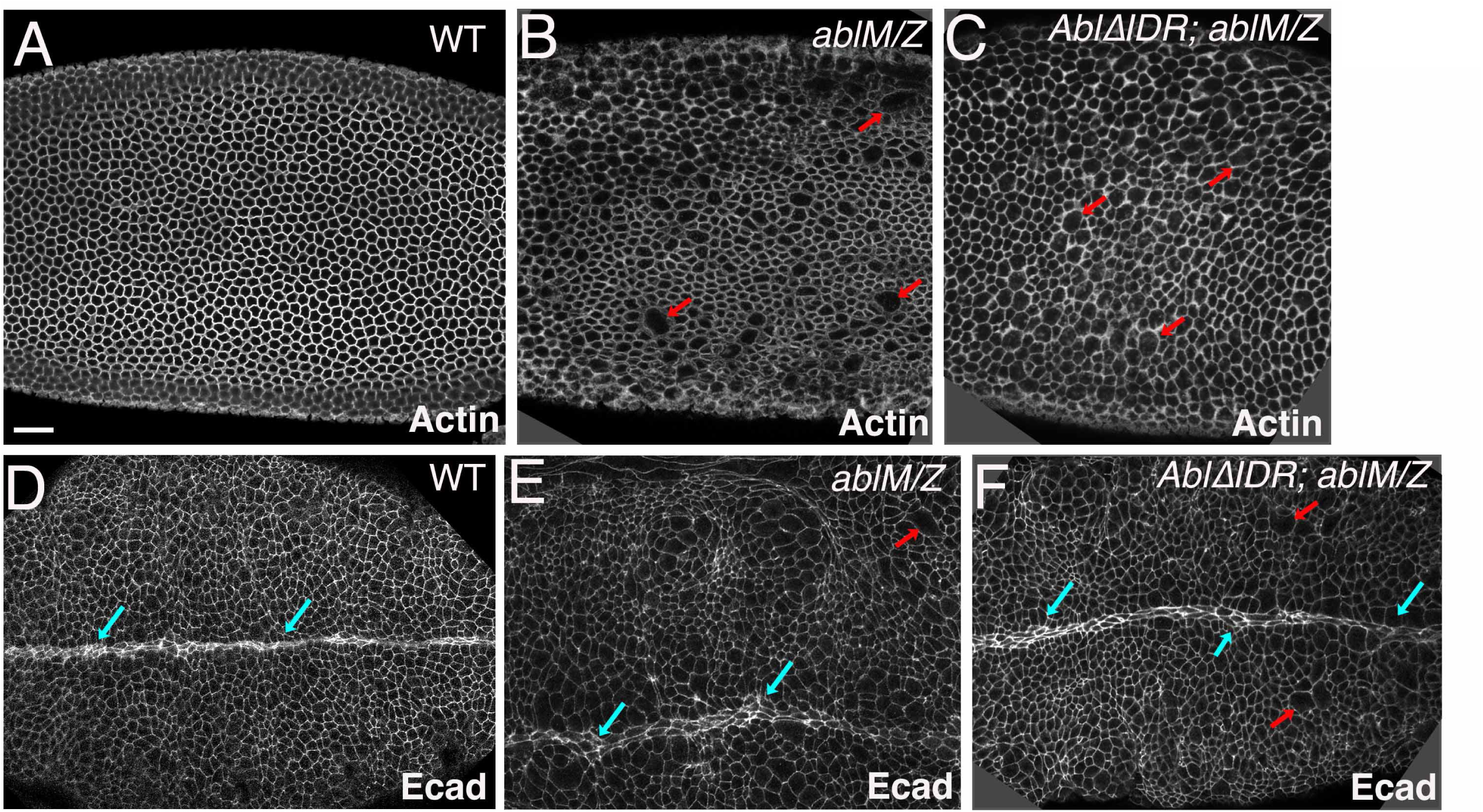
AblΔIDR does not effectively rescue defects in cellularization or mesoderm invagination. Embryos, genotypes indicated, anterior left. A-C. Cellularization, Phalloidin stained to reveal f-actin. A. Wildtype. Cellularization was completed normally, producing solely mononucleate cells. B. *ablMZ* mutant. Defects in actin regulation during syncytial development and cellularization led to the formation of multinucleate cells (red arrows). C. *AblΔIDR; ablMZ* mutant. AblΔIDR fails to rescue the defect in cellularization, and thus multiple multinucleate cells are observed. D-F. Stage 8 embryos, stained with antibodies to Ecad to visualize cell shapes. D. In wildtype mesoderm invagination is completed normally leaving a straight and even midline (blue arrows). E. *ablMZ* mutant. Defects in mesoderm invagination leave the ventral midline wavy and uneven (blue arrows). Also note the continued presence of multinucleate cells (red arrows). F. *AblΔIDR; ablMZ* mutant. AblΔIDR fails to fully rescue the defect in mesoderm invagination, leaving a wavy midline (blue arrows). Multinucleate cells remain (red arrows). Scale bar=15 µm.

### AblΔIDR does not rescue defects in germband retraction or dorsal closure

The morphogenetic events in which Abl’s roles have been analyzed in greatest detail are two of the final morphogenetic movements of embryogenesis: germband retraction and dorsal closure (Grevengoed *et al*. 2001; Rogers *et al*. 2016). These events are easily visualized by staining embryos with antibodies to E-cadherin (Ecad) to outline cells. At the end of stage 11 of wildtype embryogenesis, the caudal end of the embryo is curled up on the dorsal side. During stage 12, the germband retracts, ultimately positioning the tail end of the embryo at the posterior end of the egg, and thus leaving structures like the spiracles at the posterior end (Fig. 2A, arrow) and out of the dorsal view. At this stage, the ventral and lateral side of the embryo are enclosed in epidermis, but the dorsal side is covered by a “temporary” tissue, the amnioserosa (AS, Fig. 4A). During dorsal closure, the epidermis and the amnioserosa work in parallel to completely enclose the embryo in epidermis (reviewed in (Hayes and Solon 2017; Kiehart *et al*. 2017). Pulsatile apical constriction of the amnioserosal cells exerts force on the epidermis. In parallel, cells at the leading edge of the epidermis assemble a contractile actin cable, anchored cell-cell at leading edge tricellular junctions--this keeps the leading edge straight (LE, Fig. 4A,B blue arrows) and is important for zippering the epidermis together as the sheets meet at the canthi (Fig. 4A,B, red arrows). Actin=based protrusions from leading edge cells also aid in cell matching/alignment between the two sheets. *ablMZ* mutants have defects in both germband retraction and dorsal closure (Grevengoed *et al*. 2001; Rogers *et al*. 2016). Germband retraction is not completed and the spiracles are thus positioned dorsally (Fig. 4C, green arrow). Dorsal closure proceeds very abnormally and often fails to go to completion. The leading edge is highly wavy rather than straight (Fig. 4C, blue arrows) and zippering at the two canthi is slowed (Fig. 4C, red arrows). Tissue tearing is often observed at the border between the leading edge and amnioserosa, leaving underlying tissue exposed (Fig 4C, asterisk).

**Figure 4.**
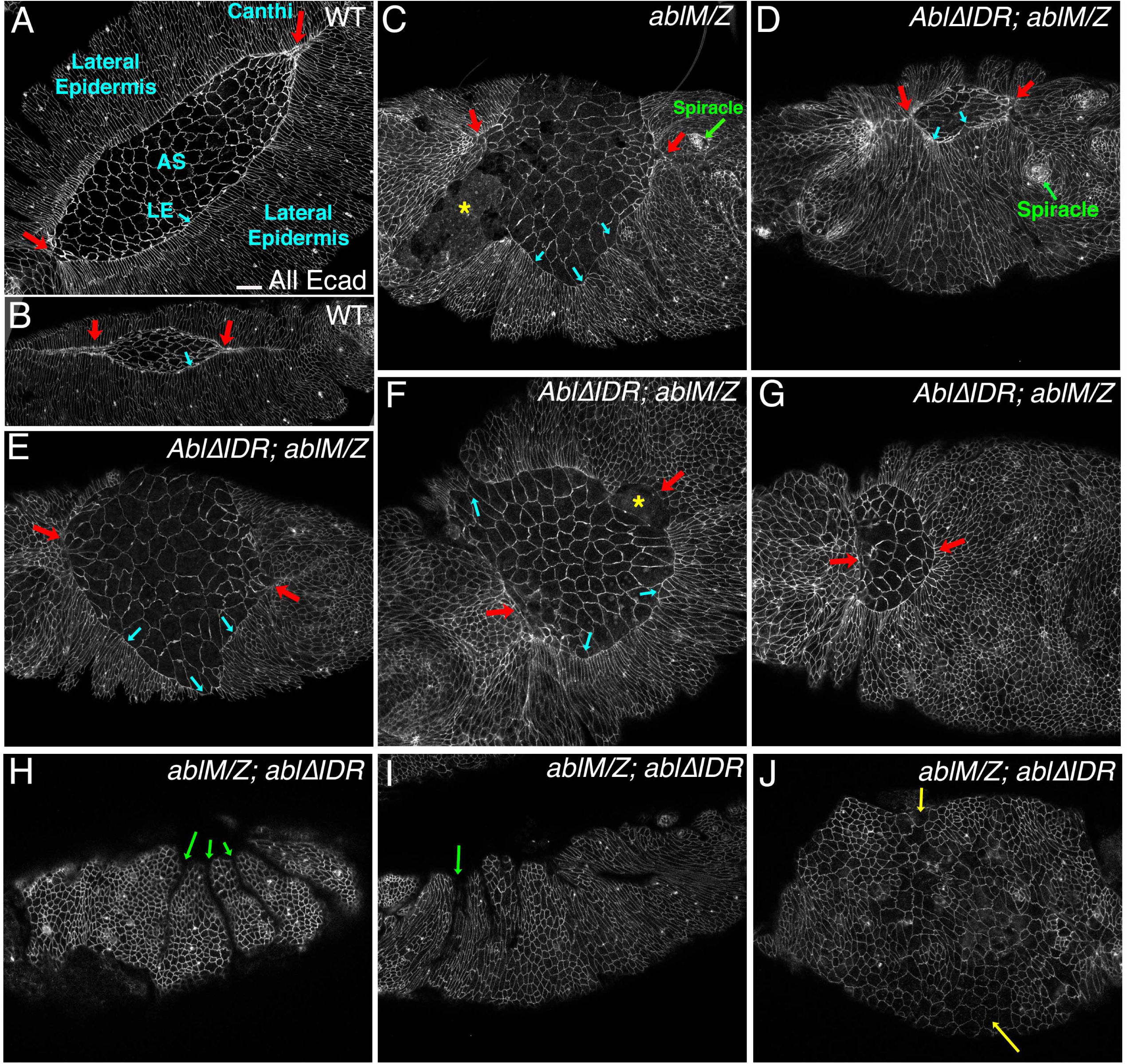
AblΔIDR does not rescue defects in germband retraction or dorsal closure. Embryos stage 13-14, anterior left, dorsal (A-G) or lateral (H-J) views, stained with antibodies to Ecad to visualize cell shapes. A,B. Wildtype embryos, dorsal view, at successively later stages of dorsal closure. The embryo is enclosed ventrally and laterally by epidermis but the dorsal surface remains covered by the amnioserosa (AS). The leading edge is straight (blue arrows) and as closure proceeds the epidermis meets and zips at the canthi (red arrows). C. Representative *ablMZ* mutant. Dorsal closure and germband retraction are disrupted. The spiracles remain dorsal (green arrow), the leading edge is wavy rather than straight (blue arrows), zipping at the canthi is slowed or halted (red arrows), and in places the amnioserosa has ripped from the leading edge, exposing underlying tissue (asterisk). D-G. *AblΔIDR; ablMZ* mutants, illustrating the range of defects in dorsal closure. D. Relatively mild phenotype, with closure nearly completed. However, the leading edge is wavy (blue arrows) and the spiracles are present dorsally, revealing failure to complete germband retraction. E,F. More typical *AblΔIDR; ablMZ* mutants, with a very wavy leading edge (blue arrows), slowed zippering at the canthi (red arrows), and ripping of the amnioserosa from the epidermis (asterisk). G. *AblΔIDR; ablMZ* mutants where zippering has happened at the posterior canthus but not the anterior one (red arrows). H-J. Most severe class of *AblΔIDR; ablMZ* mutants, in which the epidermis is reduced in extent, very deep and persistent segmental grooves remain (green arrows) and multinucleate cells are often observed (J, yellow arrows). Scale bar = 15 µm.

We thus asked whether these defects are rescued by AblΔIDR. Occasional AblΔIDR; *ablMZ* embryos succeeded in proceeding through closure, but even these exhibited defects in germband retraction, with the spiracles positioned dorsally (Fig. 4D, green arrow). In most embryos closure was highly aberrant. The leading edge was wavy instead of straight (Fig. 4D,E,F vs. A,B,blue arrows). Zippering at the canthi was slowed (Fig. 4E,F red arrows) and often did not proceed uniformly, with zippering slower or absent at the anterior end (Fig. 4G, red arrows). As we observed in unrescued *ablMZ* mutants, tearing occurred between the leading edge and the amnioserosa (Fig. 4F, asterisk). In a subset of AblΔIDR; *ablMZ* embryos, the phenotype at stage 13/14 was even more severe. These embryos had reduced epidermal coverage (Fig. 4H-J), suggesting earlier cell death. They also exhibited deep, un-retracted segmental grooves during dorsal closure (Fig. 4H,I, green arrows), another known phenotype of *ablMZ* mutants (Rogers *et al*. 2016). Some AblΔIDR; *ablMZ* embryos had numerous very large cells (Fig. 4J), which we suspect are a remnant of the multinucleate cells that arise during cellularization and gastrulation. This severe class of embryos likely represents the subset whose cuticles show substantial epithelial disruption (Fig. 2). Together, these data reveal that AblΔIDR fails to rescue defects in germband retraction or dorsal closure. Intriguingly, our previous analysis revealed that kinase activity and the FABD are largely dispensable for these morphogenetic events, while the CR1 PXXP motif in the IDR plays a role (Rogers *et al*. 2016).

### AblΔIDR does not rescue defects in leading edge cell shape or in actin regulation

We next explored the role of Abl’s IDR at the cellular and subcellular level. During dorsal closure the leading edge cells assemble a contractile actin cable that exerts tension along the dorsal cell margin. This cable maintains a straight leading edge and together with amnioserosal apical constriction elongates epidermal cells along the dorsal-ventral axis (reviewed in (Hayes and Solon 2017; Kiehart *et al*. 2017). The cable is anchored cell-to-cell at leading edge adherens junctions. In wildtype embryos tension along the cable is balanced among the cells and thus they exhibit relatively uniform shapes (Fig. 5A, arrows), with slight deviation at the segmental grooves (Fig. 5A, arrowheads). Loss of Abl disrupts leading edge cell shapes, with some cells hyper-constricted and other splayed open, presumably due to failure of the leading edge actin cable in some cells (Grevengoed *et al*. 2001; Rogers *et al*. 2016). We thus examined if AblΔIDR rescued these cell shape defects. It did not. AblΔIDR; *ablMZ* embryos exhibited penetrant defects in leading edge cell shape, with splayed open and hyperconstricted cells (Fig. 5B,C magenta vs. yellow arrows). We also observed groups of cells that failed to elongate (Fig 5B,C,E red asterisks), as we had previously observed in *ablMZ* mutants (Grevengoed *et al*. 2001; Rogers *et al*. 2016). Finally, most embryos exhibited another *ablMZ* mutant phenotype (Grevengoed *et al*. 2001; Grevengoed *et al*. 2003): large, presumably multinucleate cells, which in some embryos were very frequent (Fig. 5D, yellow asterisks; the frequency of multinucleate cells appeared substantially higher than was seen in un-rescued *ablM*Z mutants). Cell shape defects and multinucleate cells were even observed in the occasional embryos which managed to close dorsally (Fig. 5E). From these data we conclude that Abl’s IDR is essential for regulating leading edge cell shape.

**Figure 5.**
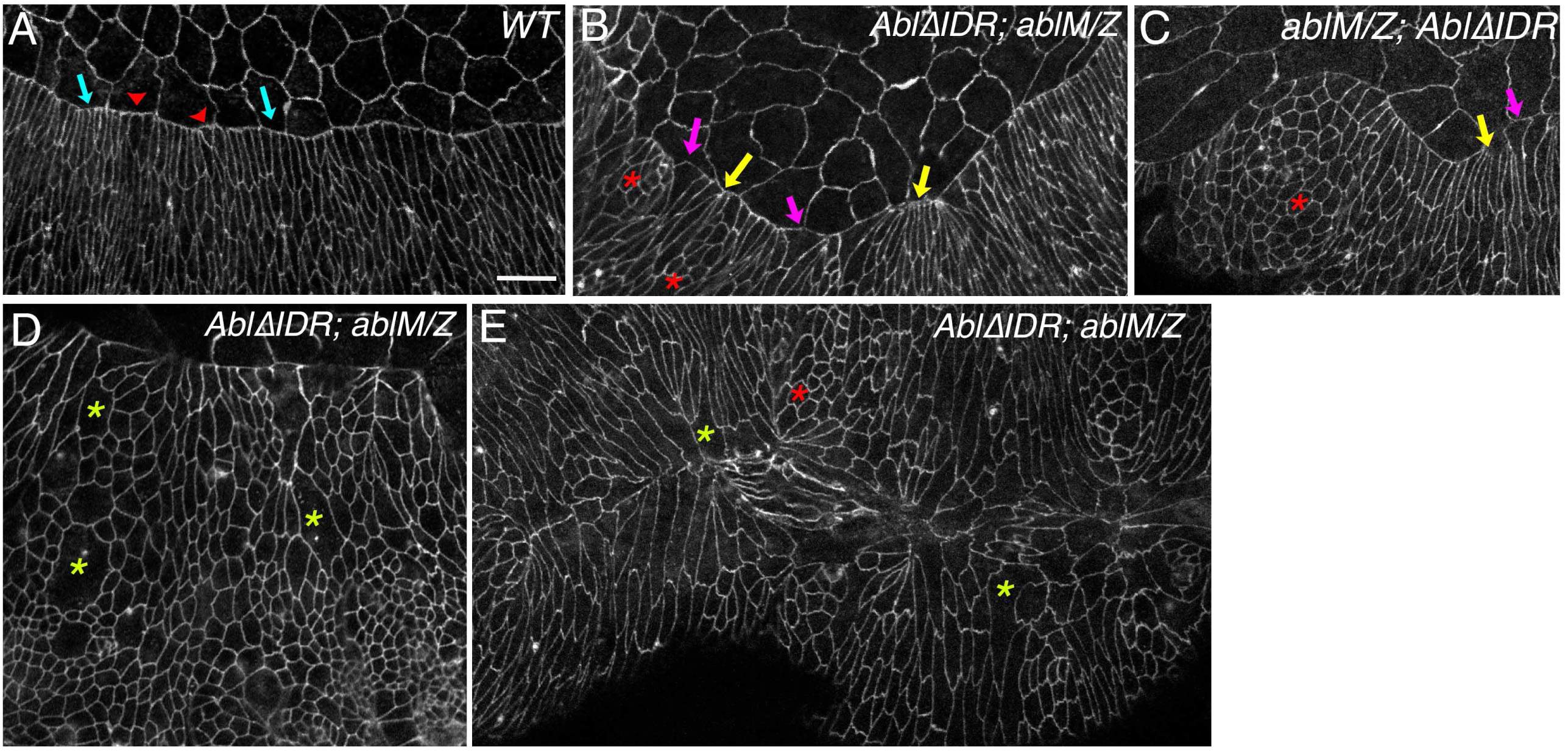
AblΔIDR does not rescue defects leading edge cell shape. Leading edge, stage 13-14 embryos, anterior left, dorsal up, stained with antibodies to Ecad to visualize cell shapes. A. Wildtype. The leading edge is straight, with even cell widths at the leading edge (blue arrows), excepting the slightly increased width at the positions of segmental grooves (red arrowheads). Scale bar=10µm. B,C. Representative *AblΔIDR; ablMZ* mutants. Leading edge cells are uneven in width, with some splayed open (magenta arrows) and some hyperconstricted (yellow arrows). Groups of cells also fail to elongate (red asterisks). D. *AblΔIDR; ablMZ* mutant. Green asterisks indicate large multinucleate cells. E. *AblΔIDR; ablMZ* mutant. Similar cell shape defects are seen in embryos that have completed or almost completed closure.

One of the key roles of Abl family kinases is regulation of the cytoskeleton. *Drosophila* Abl regulates the actin cytoskeleton through effectors like the actin polymerase Enabled (Ena). Our previous analysis suggests an important role for Abl regulation of Ena and actin at the leading edge during dorsal closure (Grevengoed *et al*. 2001; Gates *et al*. 2007; Rogers *et al*. 2016). In wildtype embryos Ena localizes to the cell junctions of both amnioserosal and epidermal cells, but is strongly enriched in the tricellular junctions of leading edge cells, where the actin cable is anchored (Fig. 6A, red arrows; (Gates *et al*. 2007; Manning *et al*. 2019). Ena is also somewhat enriched at tricellular junctions of more ventral epidermal cells (Fig. 6A, yellow arrows). In *ablMZ* mutants the uniform localization of Ena to leading edge tricellular junctions is lost (Rogers *et al*. 2016). We thus asked whether AblΔIDR can restore leading edge Ena localization. While Ena remained enriched at some leading edge tricellular junctions of AblΔIDR; *ablMZ* mutants (Fig 6B,C,D red arrows), its uniform enrichment was lost, even though enrichment at lateral epidermal tricellular junctions remained (Fig 6B,C,D yellow arrows). At many leading edge tricellular junctions Ena was weak or absent (Fig 6B,C,D cyan arrows), and at other places Ena spread across the leading edge (Fig 6B,C,D green arrows), all features we previously observed in *ablMZ* mutants and in embryos lacking the CR1 PXXP motif (Rogers *et al*. 2016).

**Figure 6.**
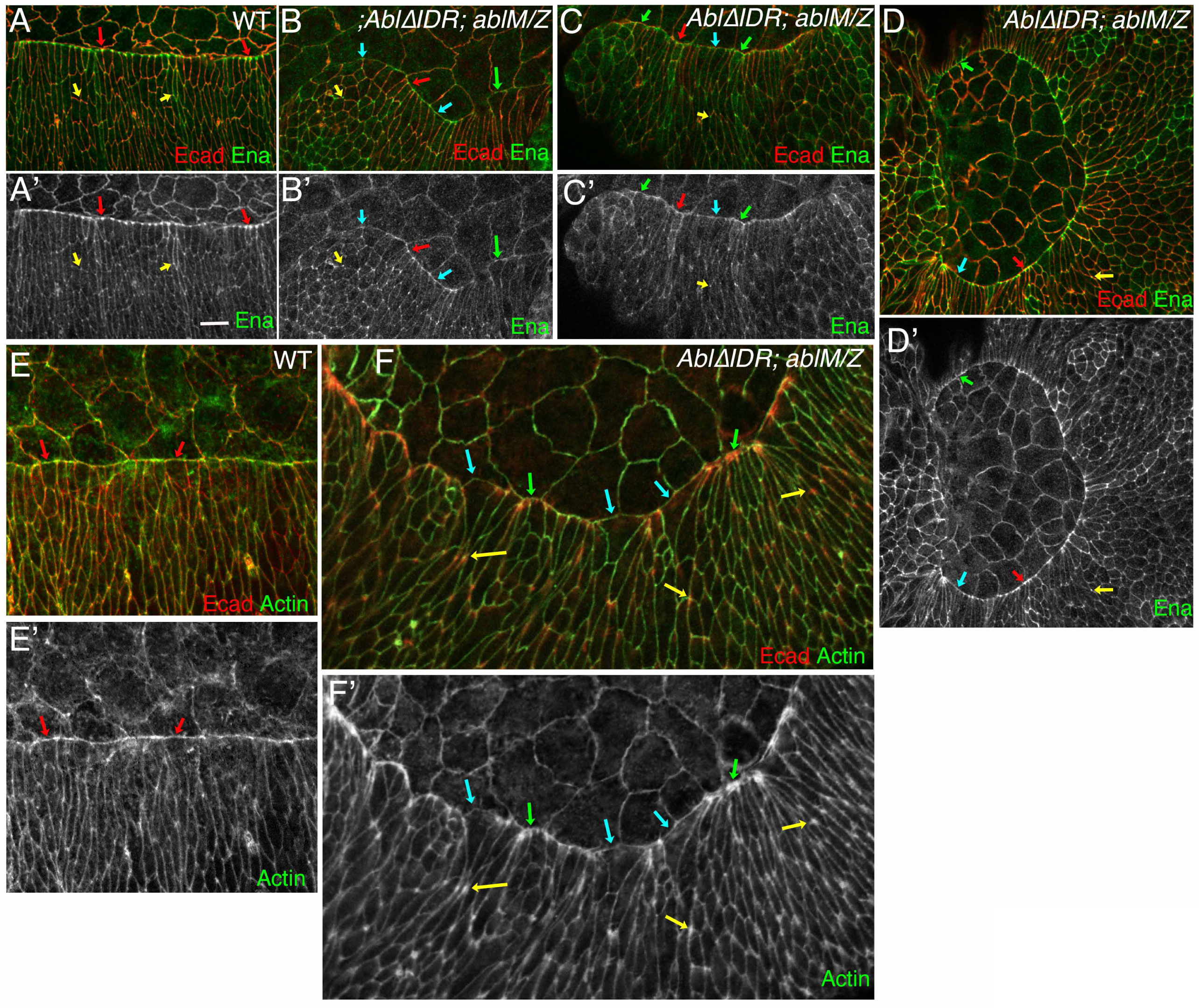
AblΔIDR does not rescue defects in Ena localization or actin regulation. Leading edge, stage 13-14 embryos, anterior left, dorsal up, stained to visualize Ecad and Ena (A-D) or Ecad and F-actin (F,G). A. Wildtype. Ena localizes cortically in both amnioserosal and epidermal cells. Ena is prominently enriched at leading edge tricellular junctions (red arrows), and is enriched at lower levels at tricellular junctions in the lateral epidermis (yellow arrows). Scale bar=10µm. B-D. *AblΔIDR; ablMZ* mutants. While Ena remains cortical and is enriched at lateral epidermal tricellular junctions (yellow arrows), uniform Ena enrichment at leading edge tricellular junctions is lost. While some tricellular junctions retain Ena enrichment (red arrows), at others Ena enrichment is reduced (cyan arrows) or Ena is found all along the leading edge (green arrows). E. Wildtype. Actin is found cortically in all epidermal cells but is enriched in the leading edge actin cable (red arrows). F. *AblΔIDR; ablMZ* mutant. While most cells still have actin along the leading edge, actin intensity varies from lower (blue arrows) to much higher than normal (green arrows). Actin is also elevated at tricellular junctions of lateral epidermal cells (yellow arrows).

The altered cell shapes observed in *ablMZ* mutants reflect defects in the leading edge actin cable (Rogers *et al*. 2016). In wildtype embryos the actin cable extends relatively uniformly across the leading edge (Fig. 6E, arrows), joined cell to cell at leading edge tricellular junctions. In contrast, in AblΔIDR; *ablMZ* embryos, the leading edge actin cable was discontinuous, with regions of reduced intensity (Fig. 6F, cyan arrows) interspersed with regions of elevated actin intensity (Fig. 6F, green arrows), as we previously observed in *ablMZ* mutants (Rogers *et al*. 2016). Actin levels were also elevated at many lateral epidermal tricellular junctions (Fig. 6F, yellow arrows) a featured shared by *ablMZ* mutants (Rogers *et al*. 2016) and by embryos in which Ena levels were artificially elevated (Nowotarski *et al*. 2014). These results indicate that AblΔIDR does not rescue the defects in Ena localization or actin regulation seen after loss of Abl.

### AblΔIDR protein is more stable than WT Abl protein

Perhaps the most surprising aspect of our phenotypic analysis of AblΔIDR were the indications that expression of this protein actually <u>enhanced</u> rather than rescued the phenotype of *ablMZ* mutants. We first examined the possibility that this reflected destabilization of the protein or a change in subcellular localization. Wildtype Abl is found in a cytoplasmic pool and is enriched at the cell cortex. Cortical enrichment is strong in early embryos and gradually reduces through the end of dorsal closure (Grevengoed *et al*. 2001; Grevengoed *et al*. 2003; Fox and Peifer 2007). Our previous analysis revealed that kinase activity and the FABD are dispensable for cortical localization, as are each of the four conserved motifs within the IDR (Rogers *et al*. 2016). To determine if there are redundant motifs in the IDR that lead to this result, we asked if AblΔIDR retained the ability to localize to the cortex. We examined this in the background of *ablMZ* mutants to eliminate the possibility of cortical recruitment via interaction with the wildtype Abl protein. At the extended germband stage endogenous Abl is enriched at the cortex, and this is mimicked by our wildtype Abl:GFP protein (Fig. 7A; (Fox and Peifer 2007; Rogers *et al*. 2016). AblΔIDR:GFP showed a similar degree of cortical enrichment at this stage (Fig. 7B,C). Cortical enrichment of both wildtype Abl:GFP and AblΔIDR:GFP was diminished during dorsal closure (Fig. 7D,E). Thus AblΔIDR encodes an apparently stable protein that retains the ability to associate with the cortex.

**Figure 7.**
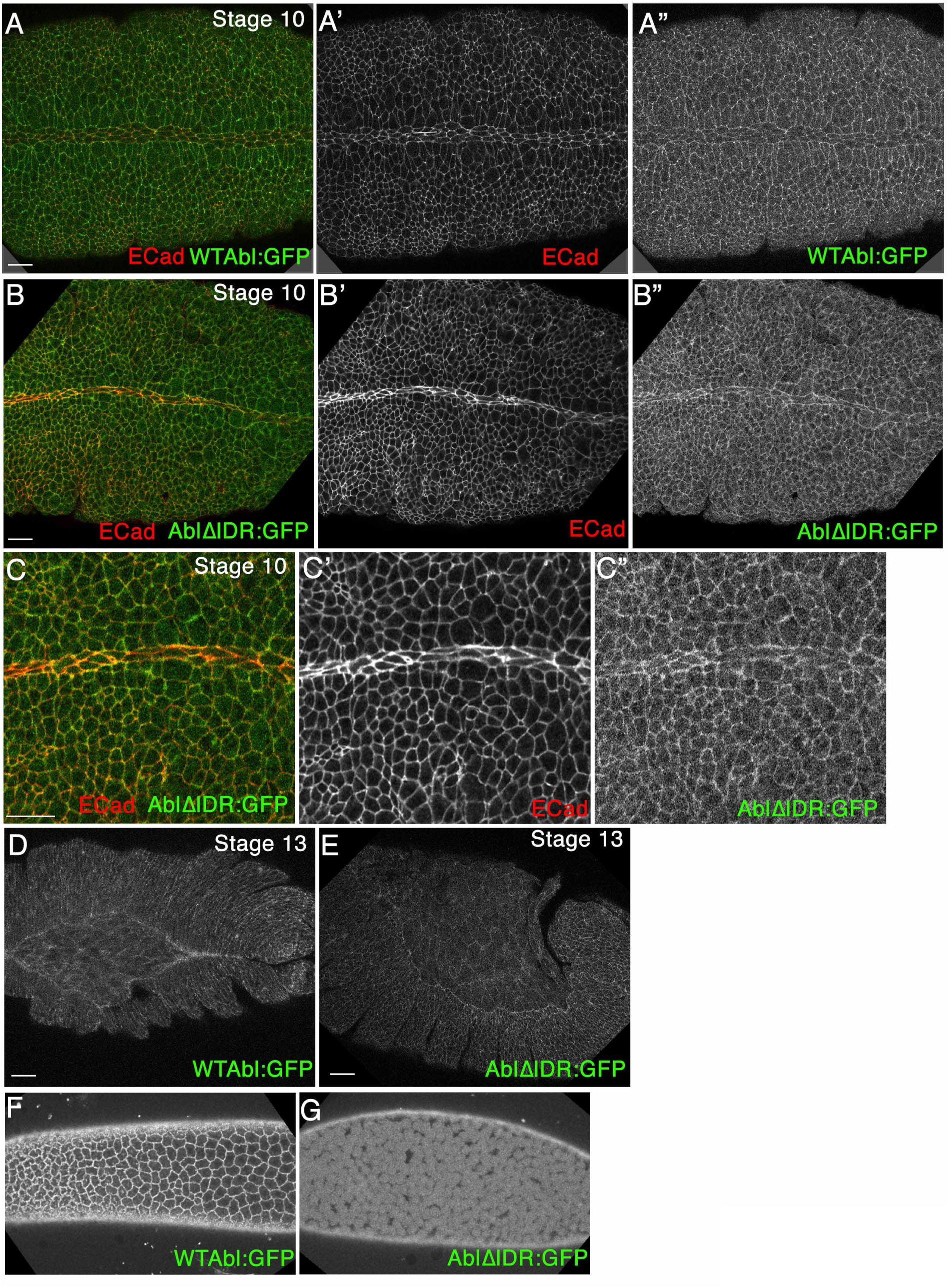
AblΔIDR:GFP protein is still enriched at the cell cortex, like wildtype Abl. A-E. Embryos, stages indicated, anterior left. Fixed and stained for Ecad, with the GFP-tagged Abl proteins directly visualized by GFP fluorescence. Scale bars=15µm. A-C. During the extended germband stage, both wildtype Abl:GFP and AblΔIDR:GFP have a cytoplasmic pool and are enriched at the cell cortex, as we previously observed is the case for endogenous Abl. D,E. Cortical enrichment of both wildtype Abl:GFP and AblΔIDR:GFP is reduced during dorsal closure. F,G. Live imaging of syncytial stage embryos. Wildtype Abl:GFP is clearly cortical but AblΔIDR:GFP is found throughout the cell.

We next visualized the Abl:GFP and AblΔIDR:GFP proteins live, without fixation. While cortical enrichment was obvious for Abl:GFP (Fig. 7F), it was much less apparent for AblΔIDR:GFP (Fig. 7G)—instead, the entire cell appeared to be filled with protein. These data suggested a second possibility: a difference in expression or accumulation levels. All of our transgenes were driven by the endogenous *abl* promotor, which drives expression of transgenes at normal levels (Fox and Peifer 2007) and in our second set of transgenes we targeted all to the same chromosomal location to reduce the possibility of position effects. Immunoblotting had previously revealed that our wildtype GFP-tagged Abl and each of our previously analyzed mutants accumulate at levels similar to endogenous wildtype Abl (Rogers *et al*. 2016). We thus repeated this analysis with AblΔIDR.

To our surprise, in embryos, AblΔIDR protein accumulates to substantially higher levels than that of wildtype GFP-tagged Abl (Fig. 8A); quantitative immunoblots revealed that protein levels are elevated 11-fold (Fig. 8B). This cannot be attributed to chromosomal position effects, as we observed similar elevation in protein levels with flies carrying two independently generated AblΔIDR transgenes (flies carrying the P-element -mediated transgenes generated for our initial experiments and the phiC targeted transgenes). Because this result was so surprising, also we expressed our transgenic proteins in a well-characterized *Drosophila* cultured cell line, S2 cells, where they were driven by the heterologous metallothionein promotor. Strikingly, AblΔIDR protein also accumulated to a significantly higher level than wildtype Abl protein in transfected S2 cells (Fig. 8C). This observation ruled out the possibility that the higher levels of AblΔIDR protein accumulation are solely due to differences in transcription, since in the embryos, transcription of both wildtype and AblΔIDR transgenes are driven by the same 2kb upstream *abl* promoter region, while in S2 cells, transcription of transgenes encoding wild type Abl and AblΔIDR was driven by the same metallothionein promoter and the plasmids encoding them had essentially identical transfection efficiencies. Consistent with what we observed in live embryos, the enrichment of Abl:GFP in the S2 cell lamellipodium was obscured when we examined AblΔIDR (Fig. 8D), consistent with the possibility that its elevated levels saturated normal binding sites in the lamellipodium and filled the cell. These data suggest that AblΔIDR protein is more stable and resistant to degradation than wildtype Abl, and that Abl’s IDR contains an element important for regulating Abl protein levels.

**Figure 8.**
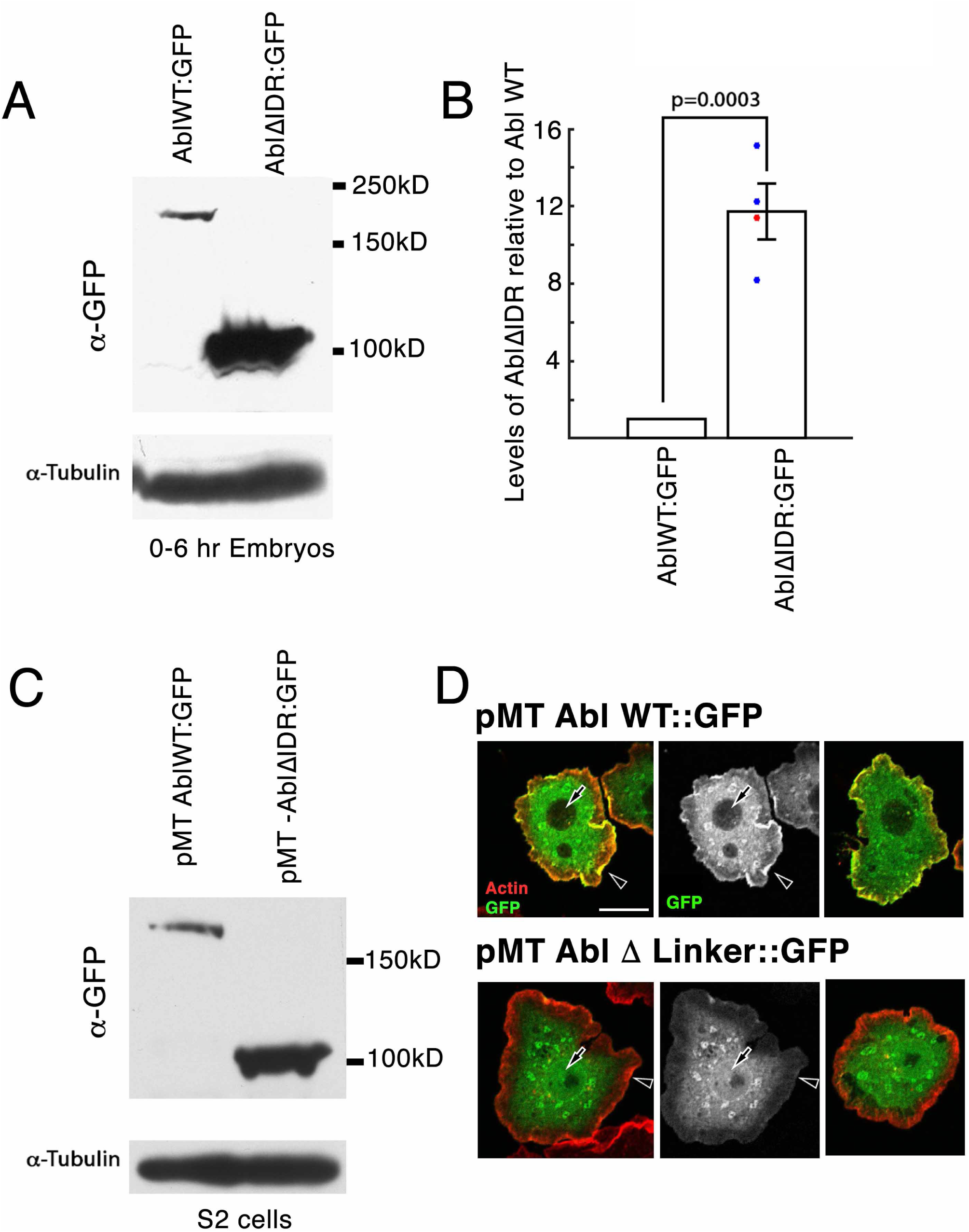
AblΔIDR protein accumulates at much higher levels than wildtype Abl. A. Immunoblot of 0-6 hr embryonic extracts, blotted with antibody to GFP to detect our transgenic proteins. Tubulin serves as a loading control. Despite the fact that both transgenes are driven by the same endogenous *abl* promotor, AblΔIDR protein accumulates at much higher levels than wildtype Abl. B. Quantification of mean protein levels from four immunoblots, normalized to both wildtype Abl:GFP and using the loading controls. Colored dots indicate values of the individual blots (Values: 8.2, 11.4, 12.3, and 15.2, Mean: 11.7; Red dot indicates blot shown in A). Error bar = standard error of the mean. C. Immunoblot of extracts of Drosophila S2 cells expressing transgenes encoding wildtype Abl:GFP or AblΔIDR, both under control of the metallothionine promotor, blotted with antibody to GFP to detect our transgenic proteins. D. Representative images of transfected S2 cells stained to visualize F-actin and our transgenic Abl proteins. Wildtype Abl:GFP is enriched in the lamellipodium (arrowhead; highlighted by F-actin) and excluded from nuclei (arrow), while AblΔIDR:GFP is not enriched in the lamellipodium or excluded from nuclei. Scale Bar=10µm.

## Discussion

The important roles of Abl kinase in embryonic development, the nervous system, adult homeostasis and oncogenesis make understanding its molecular function essential for both basic scientists and clinicians. Abl is a complex multidomain protein and we and others have assessed the roles of kinase activity and its many protein interaction domains. Here we further explore the roles of its intrinsically disordered linker (IDR), revealing this region to be essential for protein function in Drosophila morphogenesis, and important in regulating protein stability.

As one of the first protein kinases implicated in cancer, attention initially focused on Abl’s kinase activity. This clearly is critical for function of the Bcr-Abl fusion protein found in chronic myeloid and acute lymphocytic leukemia, and drugs targeting kinase activity revolutionized treatment of these disorders (Sawyers *et al*. 2002; Talpaz *et al*. 2002). However, studies of Abl’s normal roles in both Drosophila and in mammals suggest kinase activity, while important, is not essential, as proteins lacking kinase activity retain residual function in vivo (Henkemeyer *et al*. 1990; Miller *et al*. 2004; Rogers *et al*. 2016). In a similar fashion, the C-terminal f-actin binding domain and other cytoskeletal interaction motifs serve important functions in some contexts, but are not essential for protein function in others (Miller *et al*. 2004; O’donnell and Bashaw 2013; Rogers *et al*. 2016; Cheong and Vanberkum 2017).

Abl’s IDR is an interesting but poorly understood feature of Abl. IDRs are found in diverse proteins and have attracted increasing interest as regions mediating multivalent interactions, thus playing a role in some cases in phase transitions leading to the assembly of biomolecular condensates (Banani *et al*. 2017; Holehouse and Pappu 2018). They contain regions of low-complexity sequence that are not well conserved, which mediate relatively non-specific interactions. IDRs also can contain short conserved motifs that mediate specific protein interactions, as is the case in Abl. Our previous analysis focused on four motifs that are well conserved among different insects, which we referred to as CR1 to CR4 (Rogers *et al*. 2016). To our surprise, three of these, including a putative consensus binding site for the Abl effector Ena, are dispensable for rescuing viability and fertility (Rogers *et al*. 2016). However, the PXXP motif embedded in CR1 proved important for function—AblΔCR1 mutants exhibited reduced adult and embryonic viability and had defects in most but not all aspects of Abl function during embryonic morphogenesis. Cheong and VanBerkum similarly found important functions for this motif in supporting adult viability and embryonic axon guidance (Cheong and Vanberkum 2017). However, the data from both groups reveal that AblΔCR1 retains residual function. Cheong and VanBerkum extended this analysis by deleting larger regions of the IDR, singly and in combination. These data further support the idea that the PXXP motif is the only <u>individually essential</u> region of the IDR. However, their gain-of-function assays and analysis of effects on protein localization suggest that the region containing the Ena-binding motif also contributes to axon localization and subtly to function.

Here we cleanly deleted the IDR while leaving the FABD intact, allowing us to directly determine whether other regions of the IDR have additional functions. Our new data strongly support this idea. In our assays of embryonic morphogenetic events in which Abl has a known role, AblΔIDR failed to rescue mesoderm invagination and maintenance of epidermal integrity, whereas AblΔCR1, lacking only the PXXP motif, retained full or substantial function (Rogers *et al*. 2016). Consistent with our data, Cheong and VanBerkum found that deleting the first quarter of the IDR had stronger effects than simply mutating the PXXP motif (Cheong and Vanberkum 2017; Cheong *et al*. 2020). In fact, loss of the IDR reduced Abl function more substantially than <u>any</u> of our other previous alterations, including simultaneously eliminating kinase activity and the FABD (Rogers *et al*. 2016), demonstrating its critical role in Abl function.

Surprisingly, in our assays of embryonic morphogenesis, expressing AblΔIDR actually <u>worsened</u> the phenotype of *ablMZ* mutants, substantially elevating the frequency of embryos with severe disruption of epidermal integrity. We similarly observed drastic disruption of epidermal integrity when we used the AblΔIDR transgene to rescue the progeny of *abl*^*4*^*/Df* females, once again a phenotype more severe than that of *ablMZ* mutants. The disruption of epidermal integrity we observe is likely a consequence of the early defects in syncytial development and cellularization, leading to the formation of multinucleate cells. Other mutants, including those that disrupt syncytial development and cellularization in different ways, as is seen in embryos mutant for the septin *peanut*, lead to a similarly disrupted cuticle phenotype (Adam *et al*. 2000). Intriguingly, embryos maternally and zygotically mutant for the adapter protein Crk, which in mammals can bind the Abl PXXP motif (Hossain *et al*. 2012; Gregor *et al*. 2019), also have strong defects in syncytial development and cellularization, leading to strong disruption of epithelial integrity (Spracklen *et al*. 2019), as we observed here. Crk regulates actin dynamics in the early Drosophila embryo by recruiting SCAR to the cortex (Spracklen *et al*. 2019), and the PXXP motif in Abl’s IDR can bind proteins in the WAVE regulatory complex (Cheong *et al*. 2020), of which Scar is a part. Together these data support an important role for the IDR in mediating Abl’s regulation of actin.

What accounts for these seemingly “dominant negative” effects of AblΔIDR? In our view, there are several possibilities, which are not mutually exclusive. First, it is possible that the allele we use as an *abl* null allele (Fox and Peifer 2007), *abl*^*4*^, actually encodes a very low levels of partially functional protein, via readthrough of the stop codon or a low level of downstream re-start. This could explain why the phenotype of the progeny of AblΔIDR; *abl*^*4*^*/Df* females is even worse than the phenotype of AblΔIDR; *ablMZ* mutants. In this scenario, AblΔIDR could interfere with the function of this residual Abl protein by forming inactive complexes with it or with some of its effectors or regulators. Consistent with this, Cheong and VanBerkum saw strong dominant enhancement of mutants lacking the axon guidance modulators Frazzled and Slit when they overexpressed an Abl mutant lacking the first quarter of the IDR (Cheong and Vanberkum 2017). It also is possible that the Deficiency we used, *Df(3L)st-7 Ki*, also deletes a gene that enhances the *abl* null phenotype, as there are many known mutants that exhibit this phenotype (e.g.,(Gertler *et al*. 1989; Gertler *et al*. 1990; Forsthoefel *et al*. 2005).

The AblΔIDR protein has an additional property that may account for its dominant negative effects and which also casts light on the regulation of Abl activity: it accumulates at levels substantially higher than wildtype Abl. We observed this effect with two different transgenes driven by the endogenous *abl* promotor inserted at different chromosomal locations, and, importantly, also observed it when we expressed AblΔIDR in cultured Drosophila cells driven by a heterologous promotor. These data imply that the IDR contains sequences that regulate Abl protein stability. None of our previous deletions of conserved motifs within the IDR (CR1-CR4) affected Abl levels (Rogers *et al*. 2016), nor did the larger deletions of portions of the IDR made by Cheong and VanBerkum (Cheong and Vanberkum 2017) suggesting this effect either involves a different region of the IDR or that it is a property of the IDR as a whole.

IDRs have clearly defined roles in regulating protein stability. Almost 80% of known degrons reside in disordered regions (Guharoy *et al*. 2016), while computational predictions suggest a large fraction of ubiquitylation sites are in disordered regions (Radivojac *et al*. 2010; Pejaver *et al*. 2014). When we used the computational prediction software UbPred to identify potential ubiquitination sites within Abl (Radivojac *et al*. 2010), 19 of 24 medium and high confidence predicted ubiquitination sites were located within Abl’s IDR, and three more were in the unstructured N-terminal region (Figure 9A,B). The presence of an IDR in a protein also accelerates proteasomal degradation, and they can act as initiation sites for proteolysis (Prakash *et al*. 2004). Additionally, Ng et al. found that presence of IDRs may serve an important role in mediating ubiquitination in response to heat shock (Ng et al, 2013). Taken together, this evidence strongly suggests that Abl’s IDR may play a role in ubiquitin-mediated protein turnover as a mechanism for Abl proteostasis. It will be interesting to determine whether this is a conserved property of the IDR across the Abl family, and by what mechanism this occurs. Mammalian Abl is regulated by the ubiquitin-proteasome system (Echarri and Pendergast 2001) and Abl can be ubiquitinated by the E3 ligase Cbl (Soubeyran *et al*. 2003). Mammalian Arg is also ubiquitinated in response to oxidative stress (Cao *et al*. 2005). Future work will determine if this is a conserved property of the IDR across the Abl family, and by what mechanism this occurs.

**Figure 9.**
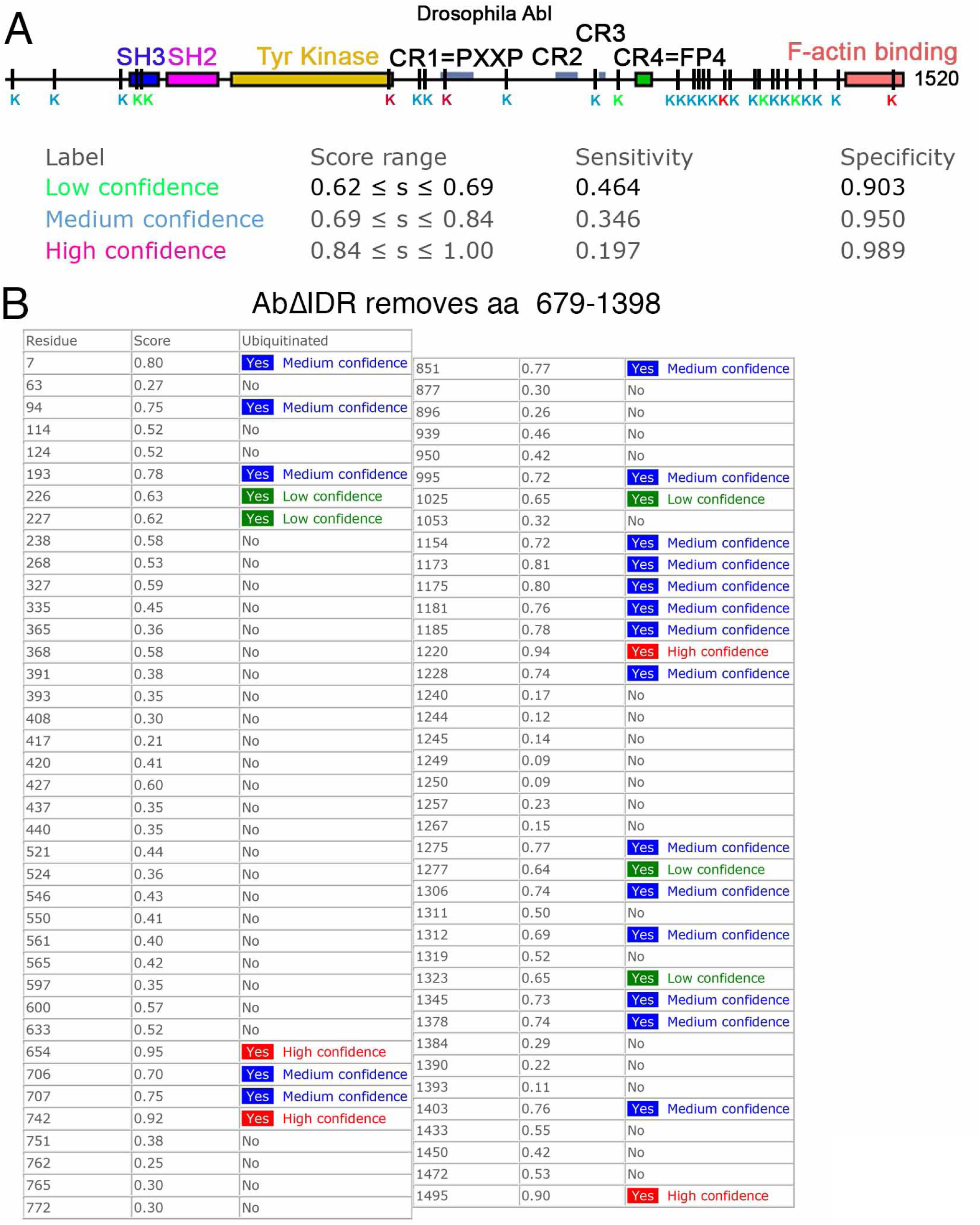
Many computationally predicted ubiqutination sites in Abl are in the IDR. Output of UbPred, computational prediction software to identify potential ubiquitination sites within Abl Abl (Radivojac *et al*. 2010). A. Diagrammatic representation, showing high (red), medium (blue) and low (green) confidence predictions. B. Table of amino acid positions of potential ubiquitination sites.

## Author contributions

E.M. Rogers and M. Peifer designed the project, E.M. Rogers and S.C. Allred carried out the experiments, E.M. Rogers, S.C. Allred and M. Peifer analyzed the data, and E.M. Rogers, S.C. Allred and M. Peifer wrote the manuscript.

## Acknowledgements

We are grateful to John Poulton for statistical advice, to Kia Perez-Vale for advice on quantifying immunoblots, to Lilia Iakoucheva for a helpful discussion of IDRs and ubiquitination, and to Steve Rogers for thoughtful comments on the manuscript. We thank the Developmental Studies Hybridoma Bank and the Bloomington Drosophila Stock Center for reagents and our lab members for thoughtful conversations. This work was supported by National Institutes of Health Grants R01 GM47957 and R35 GM118096 to M.P., and E.M.R was supported by a Leukemia and Lymphoma Society Career Development Program Fellowship Grant 5339-08.

## Materials and Methods

### Transgenic Fly Lines

To create the AblΔIDR transgene, a pair of overlapping PCR products were generated with Phusion high fidelity DNA polymerase (NEB) using pUAS-Abl:GFP (Fox and Peifer 2007) as a template. pUAS-Abl:GFP contains 2kb of 5’ upstream promoter from the endogenous *abl* gene as well as an in frame eGFP tag. The ΔIDR deletion was introduced by mutagenic DNA oligonucleotide primers in the overlapping section of the PCR products. In addition, a 15 amino acid flexible linker (GGS)_5_) was added at the location of the deletion –the hydrophilic glycine and serine residues are unlikely to form secondary structures, reducing the likelihood that the linker will interfere with the folding and function of the adjacent kinase and FABD domains. The two overlapping PCR products were joined by PCR stitching and cloned into the Xba/Not fragment of pUASg-Abl:GFP to make pUASg-AblΔIDR:GFP. The primers used for mutagenesis were as follows :

AblΔIDR Forward: 5’*GGTGGATCCGGTGGATCAGGTGGATCCGGTGGTAGTGGTGGATCC***GCCACGCCTATTGCCAAACTGA CCGAA**3’

AblΔIDR Reverse: 5’*GGATCCACCACTACCACCGGATCCACCTGATCCACCGGATCCACCG***GCTCCTCCGCCGGTGGCCACGCC CGA3**’

Italicized regions contain the code for the 15 aa flexible linker and the bold regions are complementary to the *abl* sequence. The resulting coding sequence spanning the deletion is: …TSGVATGGGAGGSGGSGGSGGSGGSATPIAKLTEP… The pUASg-AblΔIDR:GFP transgene was inserted via P-element transposition, and we were able obtain multiple independent lines, including on the 2^nd^ chromosome. To make the targeted ΔIDR transgene, the insert was excised from pUASg-AblΔIDR:GFP with Xba1 and Not1 and ligated into pUASt-attP to make pUASt-attP-AblΔIDR:GFP. The targeted transgene was targeted to the left arm of the 2nd chromosome by phiC31 integrase-mediated transgenesis into PBac{yellow[+]-attP-3B}VK00037 (cytogenetic map position: 22A3; (Bischof *et al*. 2007). Injections of transgenic constructs were performed by BestGene Inc.

### Fly Stocks, viability and phenotypic analysis of *abl* mutants and statistical tests

All experiments were done at 25°C unless noted. *y* w served as wildtype in our experiments. For assessing rescue of adult viability, we generated zygotic *abl* mutants by crossing *Df(3L)st-7 Ki/TM3 Sb* females to *transgene/transgene; abl*^*4*^*/TM3 Sb* males, and selecting for *Ki* and against *Sb* (*abl*^*4*^*/Df(3L)st-7 Ki*). We set the fraction of progeny with this genotype seen when using the wildtype *abl* transgene (AblWT; 27%) as 100%, and other genotypes were normalized to this. Adult viabilities were compared by Fisher’s Exact test (GraphPad). For this, the number of viable mutant adult flies (# of *abl*^*4*^*/Df adults)* was compared to the estimated number of non-viable flies. The number of non-viable flies was estimated by subtracting the number of viable mutant adult flies from the expected number if they were fully viable (# of *abl*^*4*^*/TM3 plus Df/TM3 divided by 2*). We used two methods to generate embryos maternally and zygotically *abl* mutant (*ablMZ*): 1) using the dominant female sterile method (Chou and Perrimon, 1996) to make *abl*^*4*^ clones in the female germline and 2) using a deficiency spanning the *abl* locus transheterozygous to *abl*^*4*^. To generate *abl* germline clones, w; *Tn[Abl]/Tn[Abl];FRT 79 D-F abl*^*4*^*/ TM3* females were crossed with *hs::Flp;;FRT 79 D-F ovoD/TM3* males. 48-72 hr old progeny were heat shocked for three hours at 37° C and allowed to develop to adulthood. Virgin female progeny of the genotype *hs::FLP/+;Tn[Abl]/+; FRT 79 D-F abl4/ FRT 79 D-F ovoD* were crossed with *w*; *Tn[Abl]/Tn[Abl];FRT 79 D-F abl*^*4*^*/ TM3, twi-GAL4,UAS-EGFP* males, embryos collected from cups with apple juice/agar plates and yeast paste As one approach, we generated embryos maternally and zygotically mutant for *abl* using a deficiency: *Df(3L) st-j7, Ki/ TM6b* (Bloomington #5416, Deletes73A2-73B2). For generation of *ablMZ* mutants by this approach *w; Df(3L) st-j7, Ki/ TM3, twi-GAL4,UAS-EGFP* females were crossed with w; *Tn[Abl]/Tn[Abl];FRT 79 D-F abl*^*4*^*/ TM3* males. The resulting w; *Tn[Abl]/+;FRT 79 D-F abl*^*4*^*/ Df(3L) st-j7, Ki* females were crossed to w; *Tn[Abl]/Tn[Abl];FRT 79 D-F abl*^*4*^*/ TM3, twi-GAL4,UAS-EGFP* males and embryos collected. Assessment of embryonic lethality and preparation of embryonic cuticles were done as in Wieschaus and Nüsslein-Volhard (1986) (WIESCHAUS AND NÜSSLEIN-VOLHARD 1986). For both approaches, embryonic viabilities were compared by Fisher’s Exact test (GraphPad). To compare cuticle phenotypes of *abl*^*4*^*MZ* mutants and embryos expressing different Abl transgenes in the *abl*^*4*^*MZ* mutant background, we used Fisher’s Exact test (GraphPad). For each genotype, the number of cuticles falling into the two more severe classes (i.e. Dorsal closure failure, and Epidermal integrity defect) were grouped to a single defective category, and compared to the number of cuticles in the two less severe categories (wildtype and Strong defects in germband retraction). Similarly, cuticle scores of each mutant transgene in the *abl*^*4*^mutant background were compared to the wildtype transgene (AblWT) using the same approach.

### Embryo live imaging

Embryos from flies that homozygous for either the transgene encoding Abl WT or AblΔIDR were dechorionated in 50% bleach and mounted in halocarbon oil (series 700; Halocarbon Products, River Edge, NJ) between a gas-permeable membrane (Petriperm; Sartorius, Edgewood,NJ) and a glass coverslip and imaged in a Z-series of 1μM slices on a Zeiss LSM-5 Pascal confocal microscope.

### Immunofluorescence

To examine embryos by immunofluorescence, flies were allowed to lay eggs on apple juice/agar plates with yeast paste for times calculated to obtain embryos at the right stages. Embryos were collected, dechorionated in 50% bleach, washed in 0.1% Triton-X, and fixed in 1:1 Heptane/3.7% Formaldehyde diluted in PBS for 20 minutes at room temperature. Embryos were then devitillenized by shaking in 1:1 heptane /methanol, when prepared for phalloidin staining, hand-devitillenized. Embryos were then blocked in Blocking Solution (PBS/0.1% Triton-X/1% Normal Goat Serum) for ≥ 30 min, incubated in primary antibody diluted in Blocking Solution overnight at 4°C and washed 3X in Blocking Solution. Embryos were then incubated in secondary antibody in Blocking Solution for 2 hours at room temperature and washed 3X in Blocking Solution. Embryos were mounted on glass slides in Aquapolymount (Polysciences, Inc). Primary and secondary antibodies were: (anti-Dcad, 1:100; anti-Enabled, 1:500 (all from the Developmental Studies Hybridoma Bank and anti-mouse and anti-rat IgG Alexa Fluors 568 and 647, from Molecular Probes); some secondary antibodies were preabsorbed with fixed *y w* embryos. For F-actin staining TRITC labeled phalloidin (Sigma) was used at a dilution of 1:500 to 1:1000.

For S2 cells, resuspended cells were allowed to attach for 1 hour onto a ConcanavalinA coated glass coverslip. Cells were then fixed 10 minutes in 10% formaldehyde HL3 buffer (70 mM NaCl; 5 mM KCl; 1.5 mM CaCl2-2H2O; 20 mM; MgCl2-6H2O; 10 mM NaHCO3; 5 mM trehalose; 115 mM sucrose; 5 mM HEPES; pH 7.2) followed by four 10 minute washes in PBS with 0.1% Triton-X (PBST) and two brief washes with ddH2O. During the last PBST wash, TRITC, labeled phalloidin was added to a dilution of 1:1000. A drop of Aquapolymount was added to the coverslips, and the coverslips were mounted on pedestals of dried nail polish on a glass slide, and sealed with nail polish. Imaging of embryos and S2 cells was done on a Zeiss LSM-5 Pascal or Zeiss 710 scanning confocal microscopes. Images were processed using ZEN 2009 software. Photoshop CS6 (Adobe) was used to adjust input levels so that the signal spanned the entire output grayscale and to adjust brightness and contrast.

### Immunoblotting

Embryonic extracts for immunoblotting were prepared by resuspending embryos in an equal volume of 2X SDS-PAGE Sample buffer(100 mM Tris-Cl (pH 6.8);4% SDS; 0.2% bromophenol blue; 20% glycerol; 200 mM B-mercaptoethanol) and homogenizing with a pestle in a microfuge tube. To make S2 cell extracts 1mL of resuspended S2 cells were spun down in a microfuge tube, the media was removed, and the pellet resuspended in an equal volume of 2X SDS PAGE Sample buffer. The samples were boiled for 5 min, spun to clear debris, and 10 ul of the resulting extract run on a 7.5% SDS-PAGE gel, and transferred to a nitrocellulose membrane. To detect the transgenic GFP-tagged Abl proteins we used anti-GFP (JL-8, 1:500 or 1:1000, Clontech). Anti-αTubulin (Sigma, 1:10000) or anti-Pnut (Developmental studies Hybridoma Bank,1:30) were used as loading controls. Detection was done using HRP-conjugated anti-mouse IgG secondary antibody (Pierce, 1:50000), and the ECL plus substrate kit (Pierce).

### Quantification of Abl ΔIDR and Abl WT protein levels

Four immunoblots of embryo extracts from homozygous stocks of the targeted Abl WT and Abl ΔIDR transgenes were used to quantify relative levels of Abl WT and Abl ΔIDR proteins in the embryos. Scans of Western blot film exposures were opened and converted to grayscale images in Adobe Photoshop. The resulting image was opened in ImageJ as a JPEG and the pixels were inverted. Rectangular ROIs of the exact same dimensions, and just large enough to contain the thickest band were drawn around the Abl protein and loading control bands. An ROI was also drawn around an unexposed area of the film for background subtraction. The mean gray value (MGV) of the ROIs for Abl and loading control proteins, and background were determined. The background subtracted MGVs of the AblΔIDR and Abl WT bands were adjusted for any loading differences by dividing them by the MGVs of their background subtracted loading controls. The background and loading control adjusted AblΔIDR and Abl WT levels were expressed as a ratio of AblΔIDR/Abl WT, normalized to the level of Abl WT which was assigned a value of 1. To determine statistical significance an unpaired t-test was used (GraphPad).

### Expression of Abl proteins in S2 cells

To express Abl and Abl ΔIDR proteins in S2 cells, the Abl and Abl ΔIDR coding regions were cloned by Gateway Technology(Invitrogen) into pMT, a vector for metal inducible protein synthesis via the metallothionein promoter. To make pMT Abl::GFP and pMT AblΔIDR::GFP, Phusion Polymerase was used to amplify the Abl and Abl ΔIDR coding regions using pUASt-attP-Abl:GFP and pUASt-attP-AblΔIDR:GFP as a template with the following primers:

AblGFP Gateway Forward:

5’CACCATGGGGGCTCAGCAGGGCAA 3’

AblGFP Gateway Reverse:

5’CCTGTTAAGCGCATTGGAGATCTGA3’

pMT Abl or pMT AblΔIDR::GFP DNAs were transfected into S2 cells grown in Sf-900 II SFM medium (Invitrogen) in the wells of 6 well plates (35mm) using Effectene transfection reagent(Qiagen) according to the manufacturer instructions. Six hours after the transfection, CuSO_4_ was added to 500 mM to induce expression of the transgenes. Cells were allowed to induce for 24 hrs and were used for both Western Blots and Immunufluorescence microscopy. Transfection efficiency was estimated by counting GFP positive cells on a dozen 143um X 143um fields on slides for immunofluorescence and dividing by the total number of cell (for blot in Fig 7c transfection efficiency: Abl WT= 65%(n=205) and AblΔIDR=57%(n=109)

## References

Adam, J. C., J. R. Pringle and M. Peifer, 2000 Evidence for functional differentiation among Drosophila septins in cytokinesis and cellularization. Molecular Biology of the Cell 11: 3123–3135.

Banani, S. F., H. O. Lee, A. A. Hyman and M. K. Rosen, 2017 Biomolecular condensates: organizers of cellular biochemistry. Nat Rev Mol Cell Biol 18: 285–298.

Bischof, J., R. K. Maeda, M. Hediger, F. Karch and K. Basler, 2007 An optimized transgenesis system for Drosophila using germ-line-specific phiC31 integrases. Proc Natl Acad Sci U S A 104: 3312–3317.

Bradley, W. D., and A. J. Koleske, 2009 Regulation of cell migration and morphogenesis by Abl-family kinases: emerging mechanisms and physiological contexts. Journal of Cell Science 122: 3441–3454.

Cao, C., Y. Li, Y. Leng, P. Li, Q. Ma et al., 2005 Ubiquitination and degradation of the Arg tyrosine kinase is regulated by oxidative stress. Oncogene 24: 2433–2440.

Cheong, H. S. J., M. Nona, S. B. Guerra and M. F. VanBerkum, 2020 The first quarter of the C-terminal domain of Abelson regulates the WAVE regulatory complex and Enabled in axon guidance. Neural Dev 15: 7.

Cheong, H. S. J., and M. F. A. VanBerkum, 2017 Long disordered regions of the C-terminal domain of Abelson tyrosine kinase have specific and additive functions in regulation and axon localization. PLoS One 12: e0189338.

Chislock, E. M., and A. M. Pendergast, 2013 Abl family kinases regulate endothelial barrier function in vitro and in mice. PLoS One 8: e85231.

Chislock, E. M., C. Ring and A. M. Pendergast, 2013 Abl kinases are required for vascular function, Tie2 expression, and angiopoietin-1-mediated survival. Proc Natl Acad Sci U S A 110: 12432–12437.

Chou, T.-B., E. Noll and N. Perrimon, 1993 Autosomal P[ovoD1] dominant female-sterile insertions in Drosophila and their use in generating female germ-line chimeras. Development 119: 1359–1369.

Colicelli, J., 2010 ABL tyrosine kinases: evolution of function, regulation, and specificity. Sci Signal 3: re6.

Courtemanche, N., S. M. Gifford, M. A. Simpson, T. D. Pollard and A. J. Koleske, 2015 Abl2/Abl-related gene stabilizes actin filaments, stimulates actin branching by actin-related protein 2/3 complex, and promotes actin filament severing by cofilin. J Biol Chem 290: 4038–4046.

Echarri, A., and A. M. Pendergast, 2001 Activated c-Abl is degraded by the ubiquitin-dependent proteasome pathway. Curr Biol 11: 1759–1765.

Forsthoefel, D. J., E. C. Liebl, P. A. Kolodziej and M. A. Seeger, 2005 The Abelson tyrosine kinase, the Trio GEF and Enabled interact with the Netrin receptor Frazzled in Drosophila. Development 132: 1983–1994.

Fox, D. T., and M. Peifer, 2007 Abelson kinase (Abl) and RhoGEF2 regulate actin organization during cell constriction in Drosophila. Development 134: 567–578.

Gates, J., J. P. Mahaffey, S. L. Rogers, M. Emerson, E. M. Rogers et al., 2007 Enabled plays key roles in embryonic epithelial morphogenesis in Drosophila. Development 134: 2027–2039.

Gertler, F., R. Bennett, M. Clark and F. Hoffmann, 1989 Drosophila abl tyrosine kinase in embryonic CNS axons: a role in axonogenesis is revealed through dosage-sensitive interactions with disabled. Cell 58: 103–113.

Gertler, F. B., J. S. Doctor and F. M. Hoffmann, 1990 Genetic suppression of mutations in the Drosophila abl proto-oncogene homolog. Science 248: 857–860.

Gregor, T., M. K. Bosakova, A. Nita, S. P. Abraham, B. Fafilek et al., 2019 Elucidation of protein interactions necessary for the maintenance of the BCR-ABL signaling complex. Cell Mol Life Sci.

Grevengoed, E. E., D. T. Fox, J. Gates and M. Peifer, 2003 Balancing different types of actin polymerization at distinct sites: roles for Abelson kinase and Enabled. J. Cell Biol. 163: 1267–1279.

Grevengoed, E. E., J. J. Loureiro, T. L. Jesse and M. Peifer, 2001 Abelson kinase regulates epithelial morphogenesis in Drosophila. J Cell Biol 155: 1185–1198.

Gu, J. J., N. Zhang, Y. W. He, A. J. Koleske and A. M. Pendergast, 2007 Defective T cell development and function in the absence of Abelson kinases. J Immunol 179: 7334–7343.

Guharoy, M., P. Bhowmick and P. Tompa, 2016 Design Principles Involving Protein Disorder Facilitate Specific Substrate Selection and Degradation by the Ubiquitin-Proteasome System. J Biol Chem 291: 6723–6731.

Hardin, J. D., S. Boast, P. L. Schwartzberg, G. Lee, F. W. Alt et al., 1995 Bone marrow B lymphocyte development in c-abl-deficient mice. Cell Immunol 165: 44–54.

Hardin, J. D., S. Boast, P. L. Schwartzberg, G. Lee, F. W. Alt et al., 1996 Abnormal peripheral lymphocyte function in c-abl mutant mice. Cell Immunol 172: 100–107.

Hayes, P., and J. Solon, 2017 Drosophila dorsal closure: An orchestra of forces to zip shut the embryo. Mech Dev 144: 2–10.

Henkemeyer, M., F. Gertler, W. Goodman and F. Hoffmann, 1987 The Drosophila Abelson proto-oncogene homolog: identification of mutant alleles that have pleiotropic effects late in development. Cell 51: 821–828.

Henkemeyer, M., S. West, F. Gertler and F. Hoffmann, 1990 A novel tyrosine kinase-independent function of Drosophila abl correlates with proper subcellular localization. Cell 63: 949–960.

Holehouse, A. S., and R. V. Pappu, 2018 Functional Implications of Intracellular Phase Transitions. Biochemistry 57: 2415–2423.

Hossain, S., P. M. Dubielecka, A. F. Sikorski, R. B. Birge and L. Kotula, 2012 Crk and ABI1: binary molecular switches that regulate abl tyrosine kinase and signaling to the cytoskeleton. Genes Cancer 3: 402–413.

Huang, Y., E. O. Comiskey, R. S. Dupree, S. Li, A. J. Koleske et al., 2008 The c-Abl tyrosine kinase regulates actin remodeling at the immune synapse. Blood 112: 111–119.

Kannan, R., and E. Giniger, 2017 New perspectives on the roles of Abl tyrosine kinase in axon patterning. Fly (Austin) 11: 260–270.

Khatri, A., J. Wang and A. M. Pendergast, 2016 Multifunctional Abl kinases in health and disease. J Cell Sci 129: 9–16.

Kiehart, D. P., J. M. Crawford, A. Aristotelous, S. Venakides and G. S. Edwards, 2017 Cell Sheet Morphogenesis: Dorsal Closure in Drosophila melanogaster as a Model System. Annu Rev Cell Dev Biol 33: 169–202.

Koleske, A. J., A. M. Gifford, M. L. Scott, M. Nee, R. T. Bronson et al., 1998 Essential roles for the Abl and Arg tyrosine kinases in neurulation. Neuron 21: 1259–1272.

Lee, J. K., P. T. Hallock and S. J. Burden, 2017 Abelson tyrosine-protein kinase 2 regulates myoblast proliferation and controls muscle fiber length. Elife 6.

Li, B., S. Boast, K. de los Santos, I. Schieren, M. Quiroz et al., 2000 Mice deficient in Abl are osteoporotic and have defects in osteoblast maturation. Nat Genet 24: 304–308.

Li, P., S. Banjade, H. C. Cheng, S. Kim, B. Chen et al., 2012 Phase transitions in the assembly of multivalent signalling proteins. Nature 483: 336–340.

Lin, Y. C., M. F. Yeckel and A. J. Koleske, 2013 Abl2/Arg controls dendritic spine and dendrite arbor stability via distinct cytoskeletal control pathways. J Neurosci 33: 1846–1857.

Manning, L. A., K. Z. Perez-Vale, K. N. Schaefer, M. T. Sewell and M. Peifer, 2019 The Drosophila Afadin and ZO-1 homologues Canoe and Polychaetoid act in parallel to maintain epithelial integrity when challenged by adherens junction remodeling. Mol Biol Cell 30: 1938–1960.

Miller, A. L., Y. Wang, M. S. Mooseker and A. J. Koleske, 2004 The Abl-related gene (Arg) requires its F-actin-microtubule cross-linking activity to regulate lamellipodial dynamics during fibroblast adhesion. J Cell Biol 165: 407–419.

Moresco, E. M., S. Donaldson, A. Williamson and A. J. Koleske, 2005 Integrin-mediated dendrite branch maintenance requires Abelson (Abl) family kinases. J Neurosci 25: 6105–6118.

Moresco, E. M., and A. J. Koleske, 2003 Regulation of neuronal morphogenesis and synaptic function by Abl family kinases. Curr Opin Neurobiol 13: 535–544.

Nowotarski, S. H., N. McKeon, R. J. Moser and M. Peifer, 2014 The actin regulators Enabled and Diaphanous direct distinct protrusive behaviors in different tissues during Drosophila development. Mol Biol Cell 25: 3147–3165.

O’Donnell, M. P., and G. J. Bashaw, 2013 Distinct functional domains of the Abelson tyrosine kinase control axon guidance responses to Netrin and Slit to regulate the assembly of neural circuits. Development 140: 2724–2733.

Oldfield, C. J., and A. K. Dunker, 2014 Intrinsically disordered proteins and intrinsically disordered protein regions. Annu Rev Biochem 83: 553–584.

Pejaver, V., W. L. Hsu, F. Xin, A. K. Dunker, V. N. Uversky et al., 2014 The structural and functional signatures of proteins that undergo multiple events of post-translational modification. Protein Sci 23: 1077–1093.

Prakash, S., L. Tian, K. S. Ratliff, R. E. Lehotzky and A. Matouschek, 2004 An unstructured initiation site is required for efficient proteasome-mediated degradation. Nat Struct Mol Biol 11: 830–837.

Qiu, Z., Y. Cang and S. P. Goff, 2010a Abl family tyrosine kinases are essential for basement membrane integrity and cortical lamination in the cerebellum. J Neurosci 30: 14430–14439.

Qiu, Z., Y. Cang and S. P. Goff, 2010b c-Abl tyrosine kinase regulates cardiac growth and development. Proc Natl Acad Sci U S A 107: 1136–1141.

Radivojac, P., V. Vacic, C. Haynes, R. R. Cocklin, A. Mohan et al., 2010 Identification, analysis, and prediction of protein ubiquitination sites. Proteins 78: 365–380.

Ren, R., 2005 Mechanisms of BCR-ABL in the pathogenesis of chronic myelogenous leukaemia. Nat Rev Cancer 5: 172–183.

Rogers, E. M., A. J. Spracklen, C. G. Bilancia, K. D. Sumigray, S. C. Allred et al., 2016 Abelson kinase acts as a robust, multifunctional scaffold in regulating embryonic morphogenesis. Mol Biol Cell 27: 2613–2631.

Sawyers, C. L., A. Hochhaus, E. Feldman, J. M. Goldman, C. B. Miller et al., 2002 Imatinib induces hematologic and cytogenetic responses in patients with chronic myelogenous leukemia in myeloid blast crisis: results of a phase II study. Blood 99: 3530–3539.

Schwartzberg, P. L., E. J. Robertson and S. P. Goff, 1991 A Substitution Mutation in the C-Abl Gene Introduced into the Murine Germ Line by Targeted Gene Disruption in Embryonic Stem-Cells. Human Gene Transfer 219: 217–226.

Soubeyran, P., A. Barac, I. Szymkiewicz and I. Dikic, 2003 Cbl-ArgBP2 complex mediates ubiquitination and degradation of c-Abl. Biochem J 370: 29–34.

Spracklen, A. J., E. M. Thornton-Kolbe, A. N. Bonner, A. Florea, P. J. Compton et al., 2019 The Crk adapter protein is essential for Drosophila embryogenesis, where it regulates multiple actin-dependent morphogenic events. Mol Biol Cell 30: 2399–2421.

Talpaz, M., R. T. Silver, B. J. Druker, J. M. Goldman, C. Gambacorti-Passerini et al., 2002 Imatinib induces durable hematologic and cytogenetic responses in patients with accelerated phase chronic myeloid leukemia: results of a phase 2 study. Blood 99: 1928–1937.

Tamada, M., D. L. Farrell and J. A. Zallen, 2012 Abl regulates planar polarized junctional dynamics through beta-catenin tyrosine phosphorylation. Dev Cell 22: 309–319.

Tybulewicz, V. L. J., C. E. Crawford, P. K. Jackson, R. T. Bronson and R. C. Mulligan, 1991 Neonatal Lethality and Lymphopenia in Mice with a Homozygous Disruption of the C-Abl Protooncogene. Cell 65: 1153–1163.

van Rosmalen, M., M. Krom and M. Merkx, 2017 Tuning the Flexibility of Glycine-Serine Linkers To Allow Rational Design of Multidomain Proteins. Biochemistry 56: 6565–6574.

Wieschaus, E., and C. Nüsslein-Volhard, 1986 Looking at embryos, pp. 199–228 in Drosophila, A Practical Approach, edited by D. B. Roberts. IRL Press, Oxford, England.

Zipfel, P. A., W. Zhang, M. Quiroz and A. M. Pendergast, 2004 Requirement for Abl kinases in T cell receptor signaling. Curr Biol 14: 1222–1231.

